# Gene context shapes spatial organization and transcriptional heterogeneity of extrachromosomal DNA in cancer cells

**DOI:** 10.64898/2026.10.28.762523

**Authors:** Lukas Frank, Georgia Marinopoulou, René H. Medema

## Abstract

Extrachromosomal DNA (ecDNA) promotes tumor adaptability by making oncogenic driver genes variably abundant, spatially mobile, and heterogeneously expressed. Transcriptional hyperactivation and spatial mobility of ecDNA-amplified genes is associated with aberrant regulatory states that could create targetable vulnerabilities in ecDNA-driven cancers. However, how gene context shapes their nuclear positioning and aberrant transcription remains unclear. To explore spatio-transcriptional <u>behavior</u> across ecDNA-amplified genes, we developed FLAGx, a multimodal imaging method that quantitatively resolves transcriptional activity and nuclear position of individual ecDNA genes *in situ*. Surprisingly, we find that gene context affects subnuclear distribution and transcriptional deregulation of ecDNA. Further, ecDNA clustering, cell cycle regulation, and drug responses depend on gene context, while transcriptional shutdown in mitosis persists across ecDNA species, arguing against pervasive mitotic transcription as a common feature. These findings show that driver gene context shapes ecDNA behavior and establish FLAGx as a useful approach to separate gene-specific from shared ecDNA features.

## INTRODUCTION

Extrachromosomal DNA (ecDNA) is a major mechanism of gene amplification in cancer. These circular DNA molecules can carry oncogenes, drug-resistance genes, and regulatory elements, and are present across diverse tumor types^1–3^. Unlike chromosomes, ecDNA lack centromeres and segregate unevenly during cell division. This random inheritance generates large cell-to-cell differences in gene copy number, allowing rapid shifts in oncogene dosage under selective pressure^4,5^. As a result, ecDNA can fuel tumor heterogeneity and therapy resistance. Consistent with this role, ecDNA is associated with poor clinical outcome in cancer patients^1,3,6^. Thus, ecDNA does not merely increase dosage of oncogenic driver genes, but enables these genes to become variably abundant, irregularly inherited, and heterogeneously expressed.

The oncogenic impact of ecDNA depends on the ability of their driver genes to remain transcriptionally active. Several hallmark studies have shown that expression is hyperactivated when a chromosomal gene is amplified on ecDNA^1,7,8^. Alterations have been associated with this transcriptional hyperactivation, including more open chromatin^7^, reduced promoter DNA methylation^9^, locally elevated histone acetylation^10^ and a lack of transcriptional repression at the onset of mitosis^11,12^. In addition to these features, ecDNA molecules can self-associate into micrometer-sized clusters (or hubs) in the nucleus, concentrating regulatory *trans* contacts and transcriptional co-activators to amplify gene expression^13,14^. Additional studies suggest that ecDNA behavior is impacted by the cell cycle. *MYC* gene containing ecDNA, for example, was reported to relocate from the nuclear periphery toward the nuclear interior at S-phase onset^15^. Moreover, ecDNA can experience disorganized patterns of DNA replication^16^ and more frequent transcription-replication conflicts^8^. Recent work further suggests that pervasive transcription of ecDNA-borne oncogenes may contribute to their physical association with mitotic chromosomes^11^. Together, these studies suggest that the transition of a gene from a chromosome to an extrachromosomal circle can override several layers of gene regulation that normally act to restrict and coordinate transcription.

However, it remains incompletely understood which alterations in transcriptional control or spatial interactions are consequences of the extrachromosomal state and which depend on the amplified driver gene context. Certain behaviors, such as fluctuating ecDNA abundance and their non-Mendelian inheritance to daughter cells are clearly ecDNA-intrinsic^17^. However, reports of ecDNA nuclear and transcriptional behavior across various cancer models suggest emerging context-dependencies. For example, the transcription-amplifying role of ecDNA clusters remains debated^18^. Some clusters appear dispensable for driver gene transcription. Others impact transcription, but may involve distinct modes of regulatory interactions with transcriptional regulators recruited to clusters^13,14,19^.

ecDNA harbor recurrent sets of cancer driver genes but are heterogeneous with regards to their specific sequence complexity and gene regulatory architecture, often resulting from rapid evolution in the tumor context ^3,20–22^. It is therefore not widely characterized which properties of the chromosomal genes are retained after ecDNA amplification, and which are overridden by ecDNA-intrinsic organization or selection pressure. However, understanding differences between the chromosomal and extrachromosomal state of the same driver gene is essential, because shared principles can create vulnerabilities that may be broadly exploitable across ecDNA-positive cancers, while context-dependencies will require focused targeting strategies. This aspect is particularly important for drug-targeting of processes linked to ecDNA driver gene expression, such as BET bromodomain and histone acetylation pathways, which have been implicated in ecDNA contacts, transcription and chromosome tethering^10,13,23^ and could therefore have broad therapeutic relevance.

A key limitation in resolving how transcriptional and spatial control of ecDNA driver genes is defined lies in the lack of approaches that quantitatively measure ecDNA abundance, nuclear position, and transcriptional activity in the same cell. (Epi-)genomics approaches integrate information about copy number, chromatin state, and regulatory interactions, but average across heterogeneous cells and individual gene copies. Genome conformation profiling by Hi-C and related approaches precisely maps spatial ecDNA contacts^10,24,25^, but cannot resolve heterogeneity of sequence-identical ecDNA molecules and their transcriptional activity. Conventional RNA and DNA fluorescence *in situ* hybridization (FISH) can visualize ecDNA spatial distribution and transcription. However, quantifying transcriptional activity per ecDNA gene copy with spatial resolution remains challenging when RNA and DNA are imaged separately or protocols require nonlinear signal amplification.

Recent spatial multi-omics and single-molecule approaches have revealed heterogeneous ecDNA behavior and linked it to local properties of the nucleus^26,27^. Yet, said methods often entail complex experimental workflows, instrumentation, and specialized data analysis, limiting their broad application. Moreover, studies seeking to identify ecDNA-specific behavior commonly compare ecDNA-amplified genes with matched chromosomally amplified gene arrays in homogeneously staining regions (HSRs)^28^. Both represent amplified gene states in cancer that can coexist or interconvert. In contrast, models comparing ecDNA driver genes with their native, chromosomal loci are less common, but particularly informative for identifying regulatory changes associated with the emergence of ecDNA. Together, these limitations constrain our understanding of how the extrachromosomal state alters gene activity and nuclear positioning, and, vice versa, to what extent gene context shapes ecDNA behavior.

Here, we developed FLAGx (FISH-based Localization and Activity analysis of Gene expression), a versatile two-step imaging method to quantify nuclear positioning and transcriptional activity of individual ecDNA driver genes in intact cancer cells. We applied FLAGx to matched cell models carrying three functionally distinct ecDNA-amplified genes, *ABCB1*, *DHFR*, and *MYC*, or their native chromosomal counterparts. We find that transcriptional deregulation upon ecDNA amplification and ecDNA clustering propensity differ markedly by driver gene. Moreover, gene-specific cell cycle transcriptional regulation can persist on ecDNA, while all three ecDNA genes retain mitotic transcriptional shutdown. Surprisingly, ecDNA also retain nuclear distribution biases resembling those of their matched chromosomal loci. Finally, drug-based perturbation of BET bromodomain proteins and histone deacetylases reveals divergent transcriptional and spatial responses across ecDNA species. Together, our findings demonstrate that key aspects of gene context-dependent regulation persist on ecDNA and establish FLAGx a unified approach to resolve these behaviors.

## RESULTS

### Multimodal imaging resolves gene position and nascent transcription *in situ*

To map the transcriptional activity and spatial positioning of individual ecDNA driver genes *in situ*, we developed FLAGx, a FISH-based assay that visualizes nascent RNA and the corresponding gene locus in the same cell (**Figure 1A**). FLAGx uses two sequential FISH staining rounds with terminally barcoded oligonucleotide probe pools that tile intronic regions of the target genes, at more than 100 unique probes per target. The simple, barcoded probe design enables multiplexed labeling of RNA and DNA targets, while preserving signal stoichiometry for accurate quantification of gene copy and local nascent RNA abundance. Protein targets can further be co-stained by immuno-fluorescence within the RNA-FISH round.

**Figure 1.**
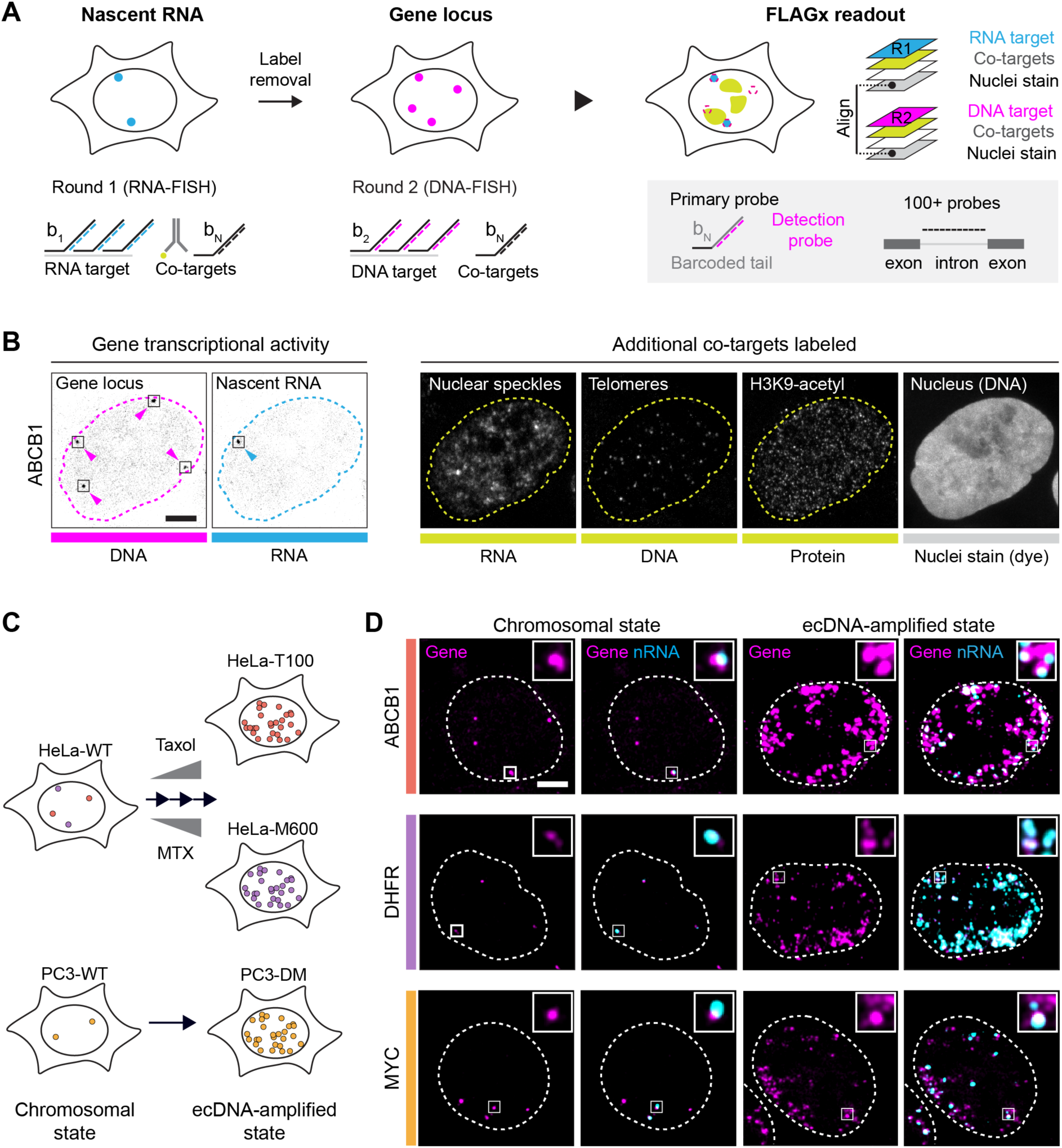
FLAGx enables integrated imaging of gene position and nascent transcription in matched ecDNA cell models. (A) Schematic of FLAGx (FISH-based Localization and Activity analysis of Gene expression). Sequential intronic RNA-and DNA-FISH detects nascent transcripts (nRNA) and corresponding gene loci using >100 unique tiling probes per target. Four fluorescent detection probes bind barcoded 3′ tails (b_N_) for multiplexed detection and linear signal amplification. Optional immunofluorescence (IF) staining detects protein targets in the first round. (B) Representative FLAGx imaging of the *ABCB1* gene in HeLa-WT cells, including nuclear speckles (*MALAT1* RNA), telomeric repeats, and H3K9ac (IF) as co-targets. (C) Scheme of background-matched chromosomal and ecDNA-amplified models of *ABCB1*, *DHFR*, and *MYC*. HeLa-T100 and HeLa-M600 are resistant to 100 nM Taxol and 600 nM methotrexate (MTX), respectively; PC3-DM is clonally derived from PC3-WT. (D) Representative FLAGx images of each cell model showing gene DNA-FISH and merged nRNA/DNA-FISH signals. Insets highlight actively transcribing chromosomal loci or ecDNA. Dotted outlines indicate nuclear boundaries. Scale bars, 5 µm.

We first benchmarked FLAGx in HeLa wild-type (WT) cells using the *ABCB1 gene* as a target locus (**Figure 1B**). FLAGx imaging resolved the chromosomal *ABCB1* loci as distinct nuclear puncta and revealed their transcriptional activity by visualizing the local presence of nascent RNA. We further demonstrated that our setup supports simultaneous detection of additional DNA, RNA, and protein targets, by co-staining of telomere repeats, *MALAT1*-RNA (nuclear speckles), and histone H3 K9-acetyl marks. RNase A digestion prior to staining largely abolished RNA signals (**Figure S1E, F)**, confirming RNA/DNA labeling specificity. Efficient removal of RNA labels before DNA locus labeling by DNA-FISH was further validated for both low-and high-abundance RNA targets (**Figure S1G, H**).

### Matched cell models link driver gene context to ecDNA state

To determine which ecDNA behaviors are shared across genes and which depend on gene context, we selected three matched cell model pairs with chromosomal and ecDNA-amplified states of the same driver genes (**Figure 1C**). We used HeLa-WT cells and two drug-resistant, ecDNA-positive derivative lines, HeLa-T100 and HeLa-M600, which amplify the *ABCB1* or *DHFR* gene on ecDNA (ec*ABCB1*, ec*DHFR*). These genes mediate resistance to the chemotherapy drugs Taxol/Paclitaxel and Methotrexate, respectively, and can be amplified and overexpressed in cancer^29–31^. Both genes differ in gene function and regulation. *DHFR* is a constitutively active, but cell cycle-regulated, metabolic gene linked to nucleotide biosynthesis. *ABCB1* encodes an ATP-dependent drug pump that is often only highly expressed upon chemotherapy resistance^32^. As third model, we selected ecDNA-negative PC3-WT prostate adenocarcinoma cells and their well-characterized isogenic derivative PC3-DM, which carries the *MYC* gene on ecDNA (ec*MYC*)^7^. *MYC* is a recurrently ecDNA-amplified oncogene encoding a growth-promoting transcription factor. To characterize these cell models, we validated the presence of ec*ABCB1*, ec*DHFR* and ec*MYC* by microscopic analysis of mitotic chromosome isolates. DNA-FISH staining (**Figure S1A-C**) alongside scoring of extrachromosomal foci on mitotic chromosome spreads (**Figure S1D**) revealed massive amplification and extrachromosomal localization of the target genes in HeLa-T100, HeLa-M600, and PC3-DM, but not in WT lines. We then established the FLAGx readout for all five cell models using probes targeting the *ABCB1*, *DHFR* and *MYC* gene sequences (**Figure 1D**). FLAGx resolved individual chromosomal loci and their nascent RNA signals in non-amplified WT cell nuclei. As expected, ecDNA models showed massively amplified gene signals, together with co-localized nascent RNA foci, indicative of ongoing transcription.

### Quantifying nuclear organization and transcription of ecDNA genes

We next leveraged our setup to extract quantitative metrics of ecDNA and chromosomal gene behavior in these models (**Figure 2A**). We first benchmarked quantification on the *ABCB1* gene in HeLa-WT and HeLa-T100 cells. To measure gene copy number on the highly variable ecDNA signals, we introduced the unit of copy number equivalent (CNE). In brief, we segmented DNA-FISH signals (gene objects) in ecDNA and WT cells, measured their intensity, and then normalized each gene object’s intensity to the average intensity of a single chromosomal locus, which was defined as 1 CNE and measured in the respective WT datasets (**Figure S2A, B**; *Methods*). This allowed us to assign a total copy number (CNE content) to nuclei, while jointly classifying ecDNA objects as singlets (CNE<2) or clusters (CNE≥2) and thereby determine the fraction of clustered ecDNA per nucleus (FClust; **Figure 2B**). Cross-validation of our CNE approach with an established, segmentation-free spot detector^33^, revealed strong agreement of both readouts across all ecDNA target genes tested (**Figure S2C-E)**. However, CNE estimation inferred more gene copies than spot detection in cells with high ecDNA abundance **(Figure S2F)**, suggesting a limitation of spot counting in dense signal clusters.

**Figure 2.**
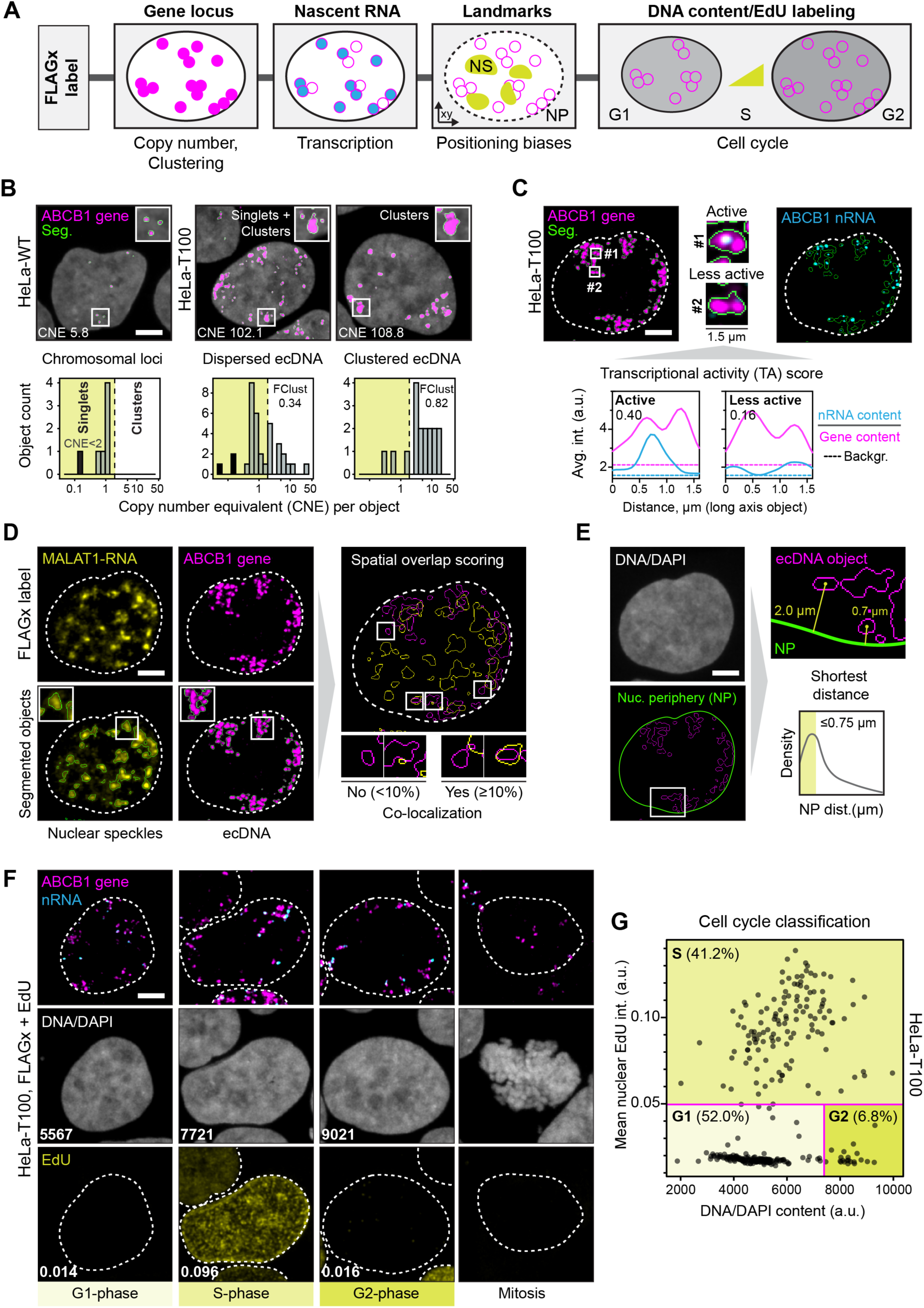
FLAGx jointly quantifies ecDNA gene copy number, clustering, nuclear positioning and transcription in single cells. (A) Overview of FLAGx-based functional readouts: gene copy number and clustering from DNA-FISH, transcriptional activity from nascent RNA enrichment, nuclear positioning relative to nuclear speckles (NS) and the nuclear periphery (NP), and cell cycle phase from DAPI/EdU staining. (B) Quantification of copy number equivalents (CNE) from segmented gene objects (green outlines) enable readout of total per cell CNE content and clustered ecDNA fraction (FClust). Chromosomal loci in WT cells define single gene copy intensity, distinguishing singlets (CNE < 2) from clusters (CNE ≥ 2). Histograms show CNE distributions for the nuclei above; black bars indicate background objects (CNE < 0.3) excluded from analysis. (C) Per-copy transcriptional activity (TA) is calculated from nascent RNA signal relative to object CNE. Insets show two ecDNA clusters with different transcriptional activities and corresponding averaged RNA/DNA intensity profiles along their long axis (1.5 µm). (D) Nuclear speckles (NS) association is defined by ≥10% overlap of gene objects (magenta) with segmented *MALAT1* RNA bodies (yellow). (E) Nuclear periphery (NP) association is defined by the distance measured from the object center to the nuclear edge (green), with ≤0.75 µm defining peripheral association. (F) Representative HeLa-T100 nuclei illustrating DAPI and EdU content (values) analysis for cell cycle classification. Mitotic cells were identified manually by morphology. (G) Classification into G1, S, and G2 by DNA/EdU content in representative HeLa-T100 cells (*n* = 311). Magenta lines indicate gating thresholds (see Methods); percentages indicate cells per phase. All scale bars, 5 µm.

To quantify per-copy transcriptional activity (TA), we divided the total nascent RNA signal associated with each gene object by its copy number equivalent (CNE; **Figure 2C**). To characterize nuclear positioning, we measured gene association with two nuclear landmarks: nuclear speckles (NS), linked to RNA splicing and active transcription^34^, and the nuclear periphery (NP), commonly associated with transcriptional repression^35^. We identified NS-associated objects by co-staining for *MALAT1* RNA^36^ (**Figure 2D**) and quantified NP proximity by measuring their shortest distance to the nuclear edge (**Figure 2E**). Finally, to investigate ecDNA behavior across cell cycle, we combined FLAGx with DNA content quantification and EdU pulse labeling to distinguish G1, S, and G2 phase cells, alongside manually identified mitotic cells (**Figure 2F, G**).

### ecDNA clustering and transcription differ markedly across driver genes

ecDNA clustering has been linked to increased transcription of ecDNA-borne genes^13,14^. We therefore asked whether clustering propensity differs across driver genes and how clustering relates to transcriptional activity (**Figure 3**). FLAGx imaging revealed clusters of ec*ABCB1*, ec*DHFR*, and ec*MYC* in cells with low, intermediate, and high ecDNA copy numbers (**Figure 3A**). However, mean ecDNA cluster size (object CNE) differed significantly between genes, being highest for ec*ABCB1* (5.8), followed by ec*MYC* (3.6) and ec*DHFR* (3.1) (**Figure 3B**).

**Figure 3.**
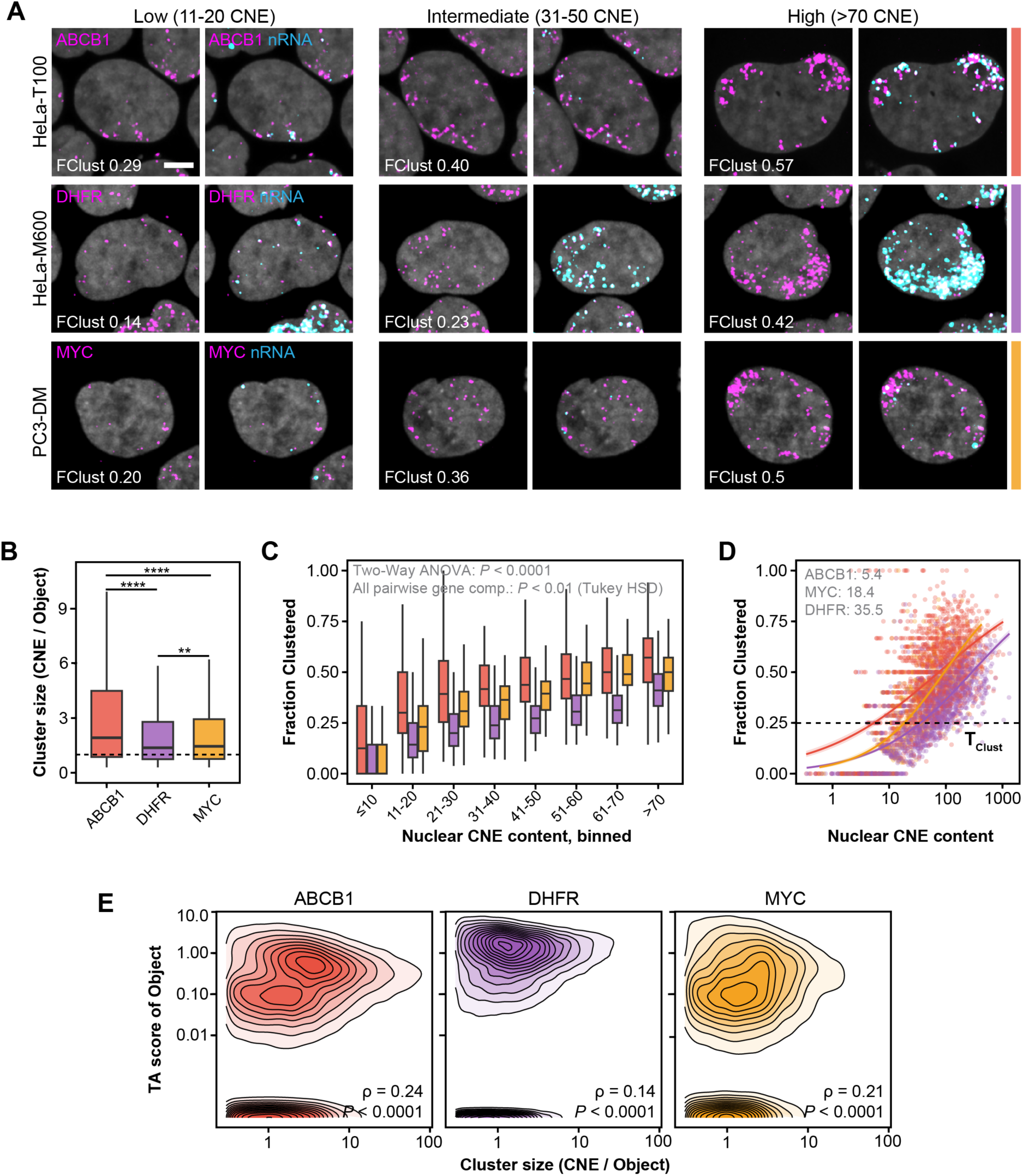
Driver gene context shapes ecDNA clustering independent of copy number. (A) Examples of FLAGx readout in HeLa-T100, HeLa-M600, and PC3-DM nuclei with low, intermediate, and high ecDNA copy number (copy number equivalent, CNE). Corresponding gene and gene/nRNA signal merges are shown with the calculated fraction of clustered ecDNA (FClust) indicated for the individual cells. Scale bars, 5 µm. (B) Distribution of mean ecDNA cluster sizes across ecDNA models. Dashed line, CNE = 1 for reference. Asterisks: pairwise Wilcoxon rank-sum tests. (C) Quantified fraction of clustered ecDNA per nucleus stratified by ranges of total copy number per nucleus (CNE content) and grouped by gene. Statistics: Two-way ANOVA with gene target and CNE bin as factors (both main effects, *P* < 0.0001; interaction, *P* = 0.00018), *P*-value for pairwise Tukey HSD post-hoc tests is indicated. (D) Clustering propensity analysis for ec*ABCB1*, ec*DHFR* and ec*MYC*. Logistic regression fit (lines with 95%CI ribbon) on CNE content versus the fraction of clustered ecDNA (dots: cells). T_Clust_, the copy number threshold at which 25% of ecDNA was clustered is indicted for each gene in units of CNE. (E) Contour plots illustrating the relationship between ecDNA cluster size and per-copy transcriptional activity (TA) score for each gene. Contour lines are equal probability density levels from 2D KDE. Rho, spearman rank correlation coefficients and *P* values (number of analyzed objects: *ABCB1*: 27,830; *DHFR*: 42,720; *MYC*: 9,672). Across panels, data comprise *N* = 2-4 independent FLAGx experiments per cell model, with *n* = 604-1681 cells per condition across experiments.

Because clustering also depends on total nuclear ecDNA copy number, we next compared the fraction of clustered ecDNA across CNE content ranges (**Figure 3C**). As expected, the clustered ecDNA fraction increased with copy number for all three genes, but significant differences remained after accounting for total copy number (Two-way ANOVA, gene effect, *P* < 0.0001; Tukey HSD, *P* < 0.01 for all gene pairs). Consistent with their mean cluster sizes, ec*ABCB1* showed the strongest clustering propensity, followed by ec*MYC* and ec*DHFR*. To further quantify these differences, we used logistic regression to estimate the nuclear copy number threshold at which 25% of ecDNA was predicted to be clustered (T_Clust_; **Figure 3D**). This threshold was lowest for ec*ABCB1* (5.1 CNE), intermediate for ec*MYC* (18.4 CNE), and highest for ec*DHFR* (35.5). Thus, ec*DHFR* required approximately 7-fold more copies than ec*ABCB1* to reach the same predicted clustering level. These results demonstrate that ecDNA clustering propensity differs markedly across genes, even after accounting for differences in copy number.

We next asked how ecDNA clustering relates to transcriptional activity. To this end, we correlated the size of individual ecDNA objects with their per-copy TA score (**Figure 3E**). Cluster size showed a weak but significant positive correlation with TA for all three genes, with varying strengths (ec*ABCB1*: ρ = 0.24; ec*MYC*: ρ = 0.21; ec*DHFR*: ρ = 0.14; *P* < 0.0001). These correlations remained significant after excluding small objects near the detection limit (**Figure S3A**). To separate the effects of cluster size and copy number, we next compared objects from cells with low and high copy numbers (bottom and top 25% of nuclear CNE content; **Figure S3B**). Across all three genes, TA increased with cluster size, but was lower overall in high-burden cells (Two-way ANOVA, both main effects, *P* < 0.0001). The burden-associated decrease varied substantially with cluster size for ec*ABCB1* and ec*MYC*, but not for ec*DHFR*. Thus, clustered ecDNA tend to be more transcriptionally active, whereas higher ecDNA copy burden is associated with reduced per-copy TA. However, the weak correlation between cluster size and TA suggests that spatial clustering alone is a poor predictor of transcriptional activity across ecDNA species.

### ecDNA amplification does not uniformly increase per-gene-copy transcription

Amplification of genes on ecDNA has been linked to increased expression and greater chromatin accessibility^7^. However, previous studies have often compared ecDNA with HSRs and measured total mRNA abundance, which does not directly reflect transcriptional activity per gene copy. This leaves unclear whether increased transcription is a general feature of ecDNA-amplified genes relative to their native chromosomal counterparts. We therefore used FLAGx to directly compare per-gene-copy transcriptional activity between matched chromosomal and ecDNA genes (**Figure 4**).

**Figure 4.**
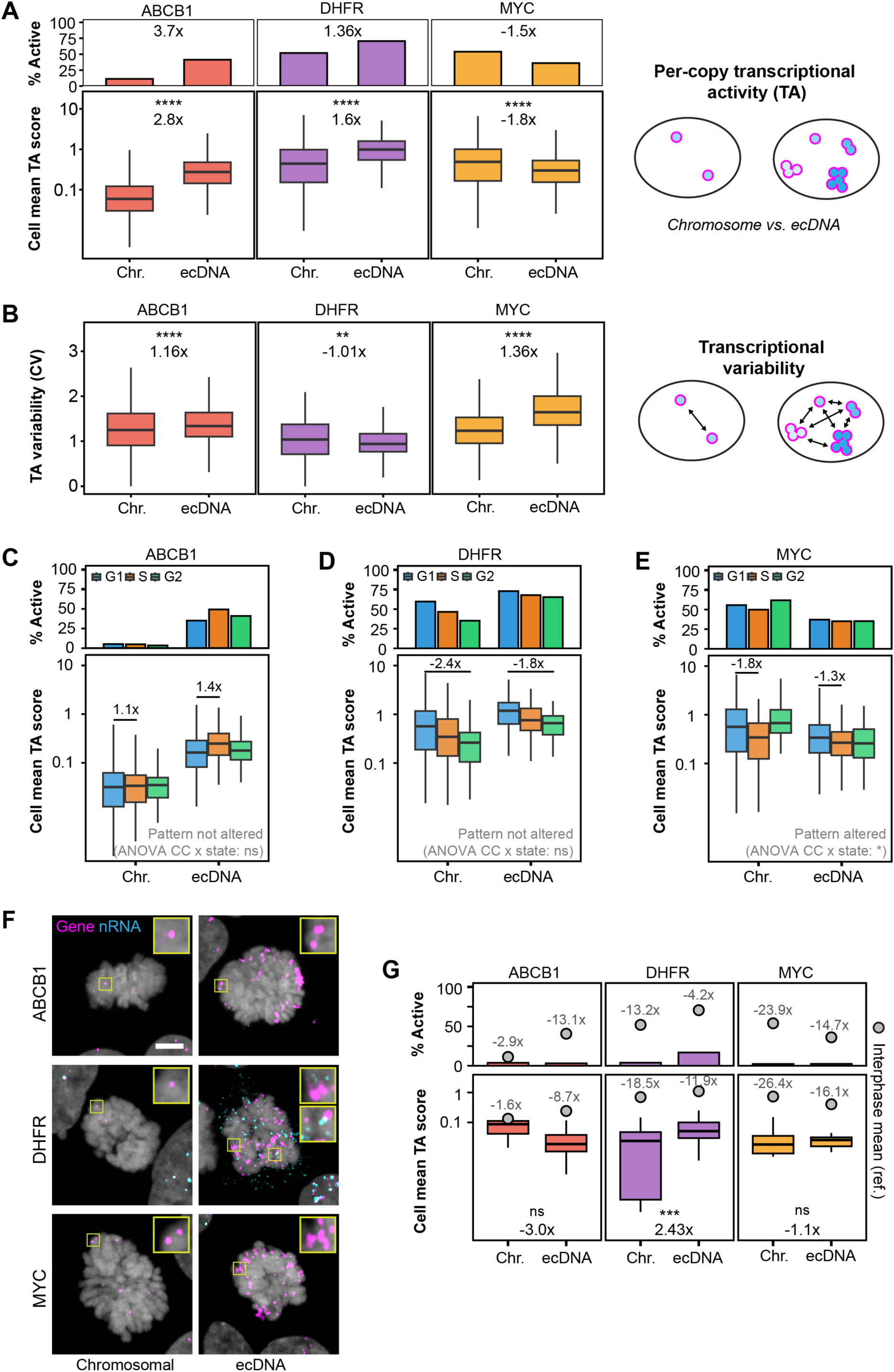
ecDNA genes retain distinct transcriptional states and cell cycle regulation. (A) Cellular means of per-copy transcriptional activity (TA; bottom) and the fraction of active gene objects (top) in matched chromosomal and ecDNA models. Fold changes relative to chromosomal means. Asterisks indicate pairwise Wilcoxon rank-sum tests. Two-way ANOVA (gene × state): main effects and interaction, *P* < 0.0001. Data comprise *N* = 2-4 independent experiments with *n* = 1191-1942 cells per condition for WT models and *n* = 604-1681 cells for ecDNA conditions (B) Within-nucleus transcriptional variability, measured as the coefficient of variation (CV) of TA among gene objects. Fold changes relative to chromosomal means; asterisks indicate Wilcoxon rank-sum tests. Dataset is identical to panel (A). (C-E) Per-cell mean TA and active fraction across G1, S, and G2 classified cells, grouped by gene state. Selected fold changes relative to G1. Two-way ANOVA (cell cycle × state) for each gene: cell cycle effects, *P* < 0.05; interactions: *ABCB1*, *P* = 0.22; *DHFR*, *P* = 0.47; *MYC*, *P* < 0.05. Data comprise *N* = 2-3 independent experiments with *n* = 856-1256 cells per condition for WT models and *n* = 604-1542 cells for ecDNA conditions. *MYC* and *DHFR* datasets are identical to those in panels (A) and (B). (F) Representative FLAGx images of mitotic cells, with enlarged chromosomal loci or ecDNA. Insets for ec*DHFR* (HeLa-M600) highlight poor colocalization of nascent RNA and DNA signals. (G) Per-cell mean TA and active fraction quantified in mitotic cells, with corresponding interphase means and relative fold changes shown in gray. One-way ANOVA with Tukey HSD tests: interphase versus mitosis, *P* < 0.001, except chromosomal *ABCB1* (*P* = 0.542); mitotic ecDNA versus chromosomal: *ABCB1*, *P* < 0.05; *DHFR*, *P* < 0.001; *MYC*, *P* = 0.092. Mitotic analyses: *N* = 2-3 independent experiments, *n* = 18-60 cells per condition. Scale bars, 5 µm.

We first quantified per-copy transcriptional activity (TA) of *ABCB1*, *DHFR*, and *MYC* across WT and ecDNA models (**Figure 4A**). In their native chromosomal state, *ABCB1* showed low transcriptional activity, *MYC* intermediate activity, and *DHFR* high activity. Upon ecDNA amplification, TA increased for *ABCB1* (2.8-fold) and *DHFR* (1.6-fold), but decreased for *MYC* (1.8-fold). An independent measure based on the fraction of transcriptionally active gene objects per nucleus showed similar trends (**Figure 4A**, top, and *Methods*). If ecDNA amplification drives a broadly hyperactive transcriptional state, individual gene copies might also become more uniformly active. We therefore asked whether amplification reduces variability in TA among gene objects within the same nucleus (**Figure 4B**). To assess this, we measured the coefficient of variation (CV) of TA among these objects. Variability changed little for *ABCB1* (1.16-fold increase) and *DHFR* (1.01-fold decrease), but increased more substantially for ec*MYC* (1.36-fold). Thus, ecDNA amplification affected both the magnitude and variability of transcription differently across genes (Two-way ANOVA, gene × state interactions, *P* < 0.0001). These findings suggest that ecDNA amplification does not uniformly increase per-copy transcription and that aspects of native, gene-specific transcriptional behavior can persist on ecDNA.

### Cell cycle transcriptional control can persist on ecDNA

Increased transcription of ecDNA-borne genes has been proposed to reflect a loss of normal gene regulatory control. We therefore asked whether ecDNA amplification preserves the transcriptional changes normally associated with the cell cycle. To address this, we used FLAGx to distinguish cells in G1, S, and G2, and mitosis (see **Figure 2F, G**). We first cross-validated our cell cycle stratification by measuring gene copy number changes across the cell cycle (**Figure S4A**). As expected, gene copy number increased significantly and by approximately two-fold from G1 to G2 across all genes, consistent with genome duplication.

We next examined how transcriptional activity (TA) changes across G1, S, and G2 in chromosomal and ecDNA states (**Figure 4C-E**). We focused on *DHFR*, whose transcription natively peaks in G1 and declines toward G2 phase^37^. In conjunction with this pattern, chromosomal *DHFR* showed highest TA in G1 and lower activity in S and G2 (1.8- to 2.4-fold reduction, **Figure 4D**). Although ec*DHFR* showed higher overall TA, it retained a similar cell cycle pattern (Two-way ANOVA, cell cycle effect, *P* < 0.0001; cycle × state interaction, *P* = 0.47). Thus, increased *DHFR* gene activity on ecDNA did not abolish its native cell cycle control. Other genes behaved differently. *ABCB1* showed a modest 1.4-fold increase in TA from G1 to S, with no significant difference in its cell cycle profile between chromosomal and ecDNA states (interaction, *P* = 0.22; **Figure 4C**). In contrast, chromosomal *MYC* showed 1.8- fold lower TA in S than G1, whereas ec*MYC* displayed a flatter profile with overall lower activity (interaction, *P* < 0.05; **Figure 4E**). Together, these findings indicate that cell cycle transcriptional regulation can persist on ecDNA, but its retention differs between genes.

Because we found larger ecDNA clusters to be associated with higher transcription (**Figure 3**), we also asked whether clustering behavior varied with cell cycle phase. All three ecDNA species formed significantly larger clusters in G2 than G1 (**Figure S4B-D**). However, total nuclear ecDNA copy number also increased from G1 to G2. We therefore tested whether the changes in cluster size could be explained by copy number (**Figure S4E**). Total copy number was strongly associated with cluster size (ANCOVA, *P* < 0.0001). After accounting for copy number, a small significant cell cycle effect remained for ec*ABCB1* and ec*DHFR* (*P* < 0.001), but not ec*MYC* (*P* = 0.19). Thus, cell cycle-associated increases in ecDNA cluster size are largely explained by higher copy number in G2 in our models, with additional minor gene-specific effects.

### Pervasive mitotic transcription is not an ecDNA-intrinsic feature

The vast majority of chromosomal genes undergo strong transcriptional shutdown during mitosis^38,39^. However, persistent mitotic transcription was found for some ecDNA-borne genes, and linked to their tethering to chromosomes^11,12^. We therefore investigated whether ecDNA is prone to escape mitotic transcriptional shutdown or whether this behavior applies only to a subset of ecDNA species. To address this, we compared transcriptional activity (TA) of *ABCB1*, *DHFR*, and *MYC* between interphase and mitosis in matched chromosomal and ecDNA models (**Figure 4F,G**). Mitotic TA was strongly reduced in both gene states and across all three genes (Tukey HSD, *P* < 0.001), with reductions reaching approximately 26-fold in mean TA and 24-fold in the fraction of active gene objects. The only exception was chromosomal *ABCB1*, which showed no significant decrease (*P* = 0.542), likely reflecting its already low interphase activity and thus limited potential for further transcriptional repression.

We next examined whether residual mitotic transcription differed between chromosomal and ecDNA states. Compared with their chromosomal counterparts, mitotic TA was 3-fold lower for ec*ABCB1* and 2.43-fold higher for ec*DHFR*, with no statistically significant difference for ec*MYC*. Despite these differences, mitotic TA remained very low compared to interphase across all ecDNA species. Notably, we detected *DHFR* nascent RNA foci in mitotic HeLa-M600 cells, which likely reflect the apparent increase in TA (**Figure 4F**). However, most of these foci did not co-localize with *DHFR* gene signals **(Figure 4F**, middle panel, insets), suggesting that the majority of these signals do not reflect ongoing transcription at the corresponding loci, but may rather be residual nascent transcripts evicted from chromatin^40^.

Together, these findings demonstrate that ec*ABCB1*, ec*DHFR*, and ec*MYC* remain largely subject to mitotic transcriptional shutdown. Thus, pervasive mitotic transcription is likely not a general feature of ecDNA, but may instead depend on the gene or cell model context.

### ecDNA retains gene-specific nuclear positioning biases

A defining feature of ecDNA is its increased spatial mobility compared with chromosomal loci, potentially exposing genes to a wider range of nuclear environments. However, chromosomal genes are not randomly positioned and their association with nuclear compartments can contribute to their transcriptional regulation. We therefore asked whether ecDNA retains gene-context specific positioning despite its greater spatial freedom and sometimes altered transcription (**Figure 5**). To this end, we measured the association of ec*ABCB1*, ec*DHFR*, and ec*MYC* with nuclear speckles (NS) and the nuclear periphery (NP) as spatial landmarks, comparing the ecDNA genes to their chromosomal counterparts (**Figure 5A**). To account for copy number effects, we grouped ecDNA-containing cells into low, intermediate, and high copy number groups. Overall, ecDNA showed higher NP association than chromosomal loci, whereas NS association was comparable (**Figure 5B,C; Figure S5A,B**). Association with both landmarks increased with copy number, but significant differences between ecDNA species remained after accounting for copy number (Two-way ANOVA, gene effect, *P* < 0.0001 for both; **Figure 5B,C**). ec*ABCB1* and ec*DHFR* showed similar NS association, but exceeding that of ec*MYC*. In contrast, NP association was strongest for ec*ABCB1*, followed by ec*MYC* and ec*DHFR* (**Figure 5C**). Strikingly, the peripheral positioning biases of the different ecDNA species closely resembled those of their matched chromosomal loci (**Figure 5D,E**). Thus, ec-DNA retains gene-specific nuclear positioning biases despite its increased spatial mobility.

**Figure 5.**
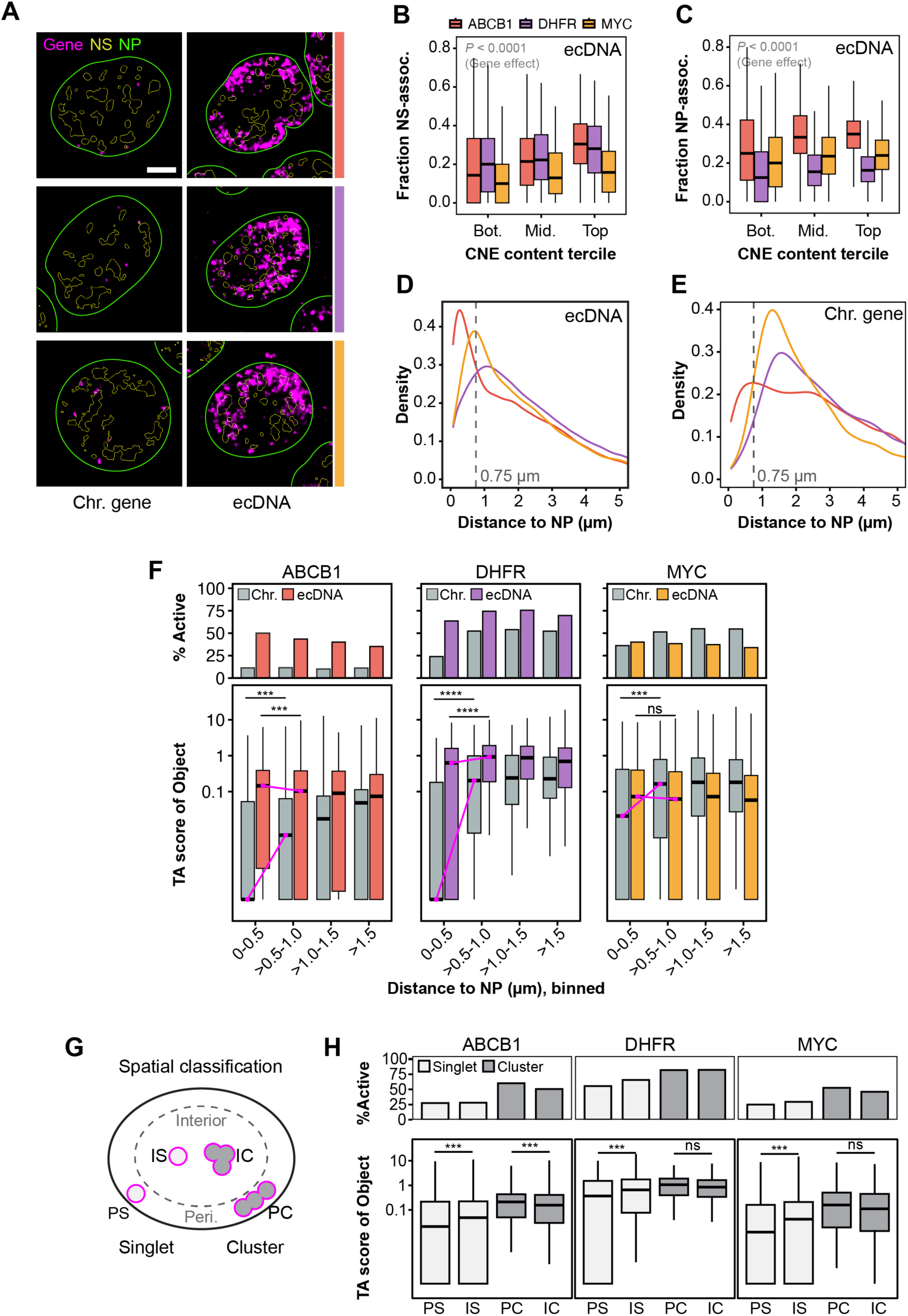
ecDNA shows gene-specific nuclear positioning and maintains transcription near the nuclear periphery. (A) Representative FLAGx images of matched chromosomal and ecDNA models, with nuclear speckle (NS) and nuclear periphery (NP) segmentation outlines. Scale bars, 5 µm. (B, C) Fraction of NS-associated (B) and NP-associated (C) ecDNA per nucleus across nuclear copy number (CNE) terciles. NP association defined as ≤0.75 µm from the nuclear edge. Two-way ANOVA (gene × CNE tercile): gene effects, *P* < 0.0001 for both. Tukey HSD: (B) ec*MYC* versus ec*ABCB1*/ec*DHFR*, *P* < 0.0001; ec*ABCB1* versus ec*DHFR*, *P* = 0.845; (C) all gene pairs, *P* < 0.0001.(D, E) NP distance distributions of ecDNA (D) and matched chromosomal loci (E), showing similar gene-specific positioning biases. (F) Per-copy transcriptional activity (TA; bottom) and active gene object fraction (top) across NP distance bins. Comparing the outermost (0-0.5 µm) and adjacent (0.5-1 µm) bins, chromosomal TA was reduced for all genes (Tukey HSD, *P* < 0.001), whereas ecDNA TA increased for ec*ABCB1* (1.5-fold), decreased for ec*DHFR* (1.3-fold), and remained unchanged for ec*MYC* (*P* = 0.996). Magenta lines connect population medians as guide to the eye. (G) Schematic classifying ecDNA as non-clustered (singlets, <2 CNE) or clustered (≥2 CNE), and as peripheral (≤0.75 µm) or internal (>0.75 µm). (H) Per-copy TA (bottom) and active fraction (top) of peripheral and internal ecDNA singlets (PS, IS) and clusters (PC, IC). Peripheral singlets showed reduced TA (Tukey HSD, *P* < 0.0001 for all genes), whereas peripheral clusters showed higher TA for ec*ABCB1* (*P* < 0.001) but no significant difference for ec*DHFR* (*P* = 0.359) or ec*MYC* (*P* = 0.692). Data across all panels comprises *N* = 2-4 independent experiments with *n* = 604-1942 cells per (gene x gene state) condition and is identical with combined datasets of Figure 4.

### ecDNA clusters retain transcriptional activity near the nuclear periphery

We next wondered how these positioning biases relate to transcriptional patterns. Consistent with the association of nuclear speckles with active transcription, NS-associated ecDNA showed significantly higher TA than non-associated ecDNA (**Figure S5C**), although the effect size was small or absent for ec*DHFR*. We then examined whether ecDNA transcription was lower near the nuclear periphery, since this environment is associated with transcriptional repression. To test this, we compared TA across 0.5 µm distance bins extending inward from the nuclear edge (**Figure 5F**). In line with prior work on transcriptional repression through nuclear lamina-association, all three chromosomal genes showed significantly reduced TA in the outermost bin (0-0.5 µm) compared with the adjacent bin (0.5-1 µm). In contrast, ecDNA showed weaker and gene-specific transcriptional responses, but largely remained at higher or equal levels in the outmost distance bin compared to the adjacent bin. Peripheral TA mildly increased for ec*ABCB1* (1.5-fold) and decreased for ec*DHFR* (1.3-fold), but remained unchanged for ec*MYC*. Thus, despite retaining gene-specific positioning biases, ecDNA transcription was overall less susceptible to peripheral repression than that of matched chromosomal loci.

This raised the question of which ecDNA remained transcriptionally active near the nuclear periphery. Given the positive association between clustering and TA (**Figure 3**), we first examined whether ecDNA clustering was biased towards the NP. Indeed, peripheral ecDNA showed increased clustering across all three genes (**Figure S5D**), with ec*ABCB1* exhibiting the strongest overall clustering propensity. To determine whether peripheral transcription differed between clustered and non-clustered ecDNA, we classified objects as singlets or clusters (**Figure 5G**) and compared TA between peripheral and internal locations within each category (**Figure 5H**). Peripheral singlets showed a moderate but consistent reduction in TA compared with internal singlets across all three genes. In contrast, peripheral clusters retained comparable TA for ec*DHFR* and ec*MYC*, and slightly higher TA for ec*ABCB1*.

Together, these findings indicate that ecDNA retains gene-specific nuclear positioning biases but is less subject to peripheral transcriptional repression than matched chromosomal loci. Although non-clustered ecDNA showed somewhat reduced transcriptional activity near the periphery, no corresponding decrease was detected for clustered ecDNA, suggesting that clustering may help maintain transcription near the repressive nuclear periphery.

### Divergent spatio-transcriptional responses of ecDNA to chromatin drugs

Identifying shared properties of ecDNA could reveal broadly targetable vulnerabilities. Previous studies suggested that BET bromodomain inhibition can disrupt BRD4-enriched ecDNA clusters and suppress transcription in ec*MYC* models^13^. Our findings revealed marked differences in clustering, transcription, and nuclear positioning across ecDNA species. We therefore asked whether perturbing chromatin-linked transcriptional regulators produces common responses across ecDNA species or whether these responses predominantly depend on gene context (**Figure 6**). To test this, we treated our models with JQ1, a BET bromodomain inhibitor, or SAHA, a histone deacetylase (HDAC) inhibitor that increases histone acetylation (**Figure 6A**).

**Figure 6.**
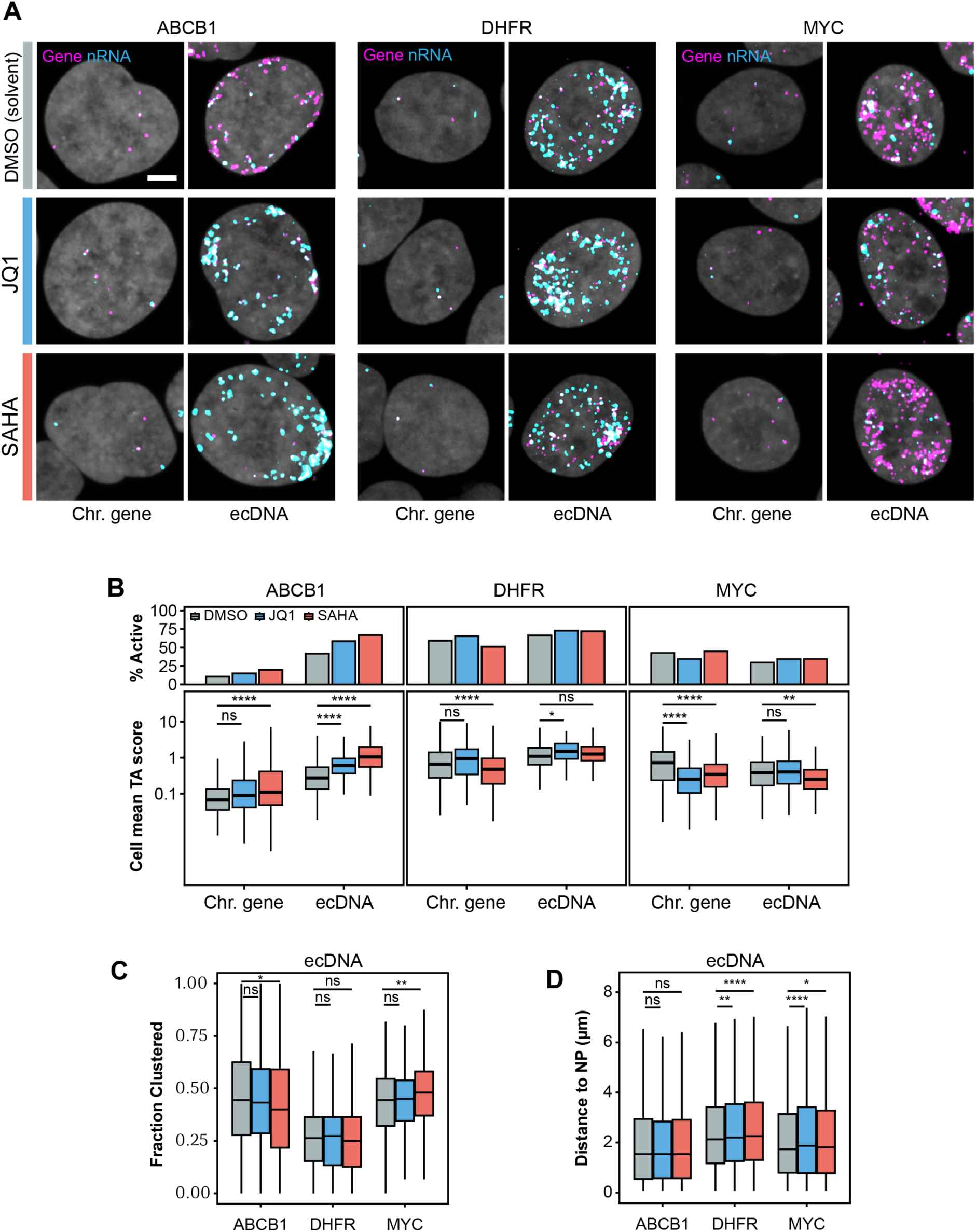
ecDNA exhibits gene-specific transcriptional and spatial responses to drugs. (A) Representative FLAGx images of matched chromosomal and ecDNA models treated for four hours with JQ1 (2 µM), SAHA (2 µM), or DMSO. Scale bars, 5 µm. (B) Per-cell mean per-copy transcriptional activity (TA; bottom) and active gene object fraction (top). Tukey HSD versus DMSO: ec*ABCB1* increased after both treatments (*P* < 0.001), ec*DHFR* increased after JQ1 (*P* = 0.015), and ec*MYC* decreased after SAHA (*P* = 0.003), but not JQ1 (*P* = 0.782). Chromosomal *ABCB1* increased and *DHFR* decreased after SAHA, whereas *MYC* decreased after both treatments (*P* < 0.01 for each). (C) Fraction of clustered ecDNA per nucleus (FClust) after drug treatments. JQ1 had no significant effect; SAHA reduced ec*ABCB1* clustering (*P* = 0.027), increased ec*MYC* clustering (*P* = 0.001), and had no significant effect on ec*DHFR* (Tukey HSD versus DMSO). (D) Distance of ecDNA objects from the nuclear periphery (NP). Both drugs induced modest inward shifts (less than 0.5 µm) for ec*DHFR* and ec*MYC* (*P* < 0.05). Data represent *N* = 2-3 independent drug treatment experiments and *n* = 464-674 cells per condition.

We first compared transcriptional activity (TA) following acute drug treatment in matched chromosomal and ecDNA models (**Figure 6B**). Responses differed markedly between genes and their ecDNA versus chromosomal states. For *ABCB1*, SAHA significantly increased TA in both chromosomal and ecDNA states, whereas JQ1 increased TA only for ec*ABCB1*. For *DHFR*, SAHA reduced chromosomal TA, while JQ1 moderately increased *DHFR* activity in its ecDNA state. Lastly, SAHA treatment significantly reduced TA of the *MYC* gene in both its chromosomal and ecDNA state, whereas JQ1 only reduced transcription for chromosomal *MYC*. Thus, drug responses differed drastically between ecDNA and matched chromosomal genes, and across driver genes. Notably, JQ1 treatment did not suppress transcription of any of the three ecDNA tested, including ec*MYC*. We next asked whether transcriptional responses to SAHA were associated with changes in histone acetylation. SAHA strongly increased nuclear H3K9ac levels across all cell models (**Figure S6A**). Notably, ec*ABCB1*, the only gene showing SAHA-dependent transcriptional upregulation, also showed a positive correlation between global H3K9ac levels and TA. No comparable association was detected for ec*DHFR* or ec*MYC* (**Figure S6B**). Thus, increased nuclear histone acetylation was not uniformly accompanied by higher ecDNA gene transcription, highlighting gene-specific responses to HDAC inhibition.

Finally, we examined whether drug treatment also altered ecDNA clustering or nuclear positioning. To our surprise, JQ1 did not significantly change the fraction of clustered ecDNA for any of the genes tested (**Figure 6C**). SAHA mildly reduced clustering of ec*ABCB1* and increased ec*MYC* clustering, but did not affect ec*DHFR*. We next assessed whether either treatment altered ecDNA positioning relative to the nuclear periphery (**Figure 6D**). Neither drug induced major spatial shifts (>0.5 µm), although ec*DHFR* and ec*MYC* showed mild biases toward the nuclear interior. Matched chromosomal loci likewise showed little spatial relocation upon perturbation, except for a small inward shift of chromosomal *MYC* with both treatments (**Figure S6C**).

Together, these findings demonstrate that acute BET and HDAC inhibition elicit distinct transcriptional and spatial responses across ecDNA species and their matched chromosomal counterparts. In particular, JQ1-induced transcriptional repression and cluster dissolution is not shared across the ecDNA species tested here, emphasizing the importance of driver gene and model context in determining drug responsiveness.

## DISCUSSION

ecDNA amplification is a major mechanism of oncogene copy number gain in cancer, contributing to tumor evolution, genetic heterogeneity, and increased oncogene expression. Several hallmark studies have established ecDNA as distinct amplification state with regulatory consequences for the genes it carries^1,2,7^. But whether ecDNA imposes common regulatory properties on amplified genes and to which extent these depend on gene context remains under investigation. Here, we developed FLAGx to jointly and directly quantify nuclear organization and transcriptional activity of ecDNA genes *in situ* at single-gene-copy resolution. Applying FLAGx to matched chromosomal and ecDNA models of *ABCB1*, *DHFR*, and *MYC* revealed that, in most cases, the ecDNA state neither uniformly increases per-copy transcription nor abolishes native regulatory control of the driver genes. Moreover, ecDNA species showed distinct clustering propensities and retained nuclear positioning biases reflecting their chromosomal counterparts, suggesting that their apparent spatial freedom remains partly constrained by gene context-specific interactions with the nuclear space. Mitotic transcriptional shutdown also persisted on ecDNA, whereas drug responses differed markedly across species. Together, these findings argue against a uniformly hyperactive and unregulated ecDNA transcriptional state.

Our direct comparison with native chromosomal loci challenges the notion that ecDNA amplification generally increases transcriptional activity per gene copy. While ec*ABCB1* showed strongly elevated per-copy transcription, ec*DHFR* was only moderately more active. Surprisingly, ec*MYC* was less active than its chromosomal counterpart. Moreover, transcriptional variability changed little for *ABCB1* and *DHFR*, suggesting that amplification does not necessarily alter the heterogeneity of transcription among gene copies. Importantly, FLAGx allowed us to distinguish increased gene copy dosage from increased transcriptional activity per copy. Our findings therefore suggest that the elevated gene expression frequently associated with ec-DNA mostly arises from increased copy number without requiring uniform transcriptional hyperactivation of the gene itself. This interpretation is consistent with recent 3D imaging-based measurements of per-copy ecDNA transcription in glioblastoma models^18^.

Our cell cycle analyses further show that native transcriptional control can persist on ecDNA. Despite increased per-copy activity, ec*DHFR* retained its characteristic decline in transcription from G1 to G2. Moreover, all three ecDNA genes underwent pronounced mitotic transcriptional shutdown, contrasting with reports of pervasive mitotic transcription in other ecDNA models^11,12^. Notably, most mitotic nascent *DHFR* RNA foci we detected in our ecDNA model did not colocalize with their corresponding gene loci, highlighting the importance of *in situ* measurements to distinguish residual RNA signals from ongoing transcription. This also raises the possibility that transcription-dependent ecDNA tethering to mitotic chromosomes may reflect regulatory interactions established before mitotic entry, rather than requiring ongoing mitotic transcription^41^. Dissecting these differences will require additional, mitotic stage-resolved measurements and mechanistic perturbations.

ecDNA clustering has been proposed to promote transcription through intermolecular regulatory contacts and the formation of BRD4-enriched transcriptional hubs^13,14^. Our findings support a positive association between clustering and transcription but show that this relationship is more complex in three respects: (i) Clustering propensity differed markedly across ecDNA species, with ec*DHFR* requiring approximately sevenfold more nuclear copies than ec*ABCB1* to reach the same predicted clustering level. (ii) Cluster size showed only a weak positive association with per-copy transcription, and correlation strength varied across genes. (iii) Although higher nuclear ecDNA copy number was overall associated with increased clustering, similarly sized ecDNA clusters were less transcriptionally active in cells with higher ecDNA burden compared to low copy cells. This could potentially reflect increased competition among ecDNA copies for limiting transcription factors or co-activators, a possibility that warrants further mechanistic investigation. Together, these findings demonstrate that clustering alone is insufficient to predict ecDNA transcriptional output and varies widely across ecDNA species. However, clustering may be particularly relevant for genes dependent on long-range enhancer-promoter interactions or, for instance, when enhancers and promoters reside on separate ecDNA circles.

Beyond clustering, the three ecDNA species we investigated also showed distinct nuclear positioning preferences. Overall, ecDNA was more frequently found near the nuclear periphery than matched chromosomal loci, but peripheral association was strongest for ec*ABCB1*, followed by ec*MYC* and ec*DHFR*, mirroring the relative positioning preferences of their native chromosomal loci. This suggests that properties of the gene locus continue to influence ec-DNA positioning. To our surprise, similar positioning did not imply similar transcription. All three chromosomal genes showed reduced transcription near the nuclear periphery, whereas their ecDNA were less consistently repressed. We also found that, across all models, ecDNA was more frequently clustered at the periphery. While peripheral non-clustered ecDNA showed reduced transcription compared with interior non-clustered ecDNA, peripheral clusters showed no corresponding reduction. Consistent with these observations, recent work reported enhanced peripheral ecDNA clustering and implicated histone H3K9me3-dependent anchoring through the lamin B receptor pathway in ecDNA positioning and gene expression^27^. Together, these findings support that clustering may help maintain ecDNA transcription near the nuclear periphery, although how clustering protects against peripheral repression remains to be resolved.

Lastly, our drug experiments revealed divergent responses across ecDNA species and their matched chromosomal loci. While JQ1 was previously shown to disrupt BRD4-enriched ec*MYC* hubs and suppress transcription in COLO320-DM cells^13^, neither effect was observed in our models, including ec*MYC* in PC3-DM. This difference may reflect the distinct BRD4-associated regulatory architecture of COLO320-DM and PC3-DM ecDNA^7^. Notably, JQ1-sensitive mitotic segregation of *MYC*-containing ecDNA has also been reported in PC3 cells^11^, suggesting that BET-dependent functions may persist even when interphase clustering and transcription are unaffected. Similarly, SAHA increased H3K9ac levels across all models, yet elicited divergent transcriptional responses, suggesting that increased histone acetylation alone does not equally affect the activity of the ecDNA gene studied here. Overall, our findings demonstrate that ecDNA amplification reshapes gene regulation without uniformly overriding the influence of gene context. Further distinguishing shared ecDNA regulatory mechanisms from those dependent on the amplified gene and cancer model background will be critical for identifying broadly targetable vulnerabilities in ecDNA-driven cancers.

## Limitations of the study

We examined three ecDNA driver genes in two cell types, limiting our ability to comprehensively separate gene-specific effects from those of ecDNA amplicon architecture and tumor context. Extending FLAGx to additional genes, including those carried on the same ecDNA molecule, and combining it with targeted genetic perturbations will help identify shared regulatory properties and dissect the causal links between clustering, nuclear positioning and transcription. We anticipate that the use of FLAGx in high-throughput screening applications will provide opportunities to comprehensively dissect these mechanistic links.

## Author contributions

L.F. and R.H.M. conceived and designed the study. L.F. performed experiments and analyzed the data. G.M. performed experiments and provided technical assistance. R.H.M. supervised the project. L.F. and R.H.M. wrote the manuscript with input from all authors.

## Materials availability

This study did not generate new unique reagents.

## Data and code availability

Custom code used in this study is made available through Zenodo, as indicated in the respective Methods sections. Microscopy data shown in the figure panels will be deposited on Figshare.

## Acknowledgements

We thank all colleagues for valuable discussions and constructive feedback, as well as the support of the Maxima Imaging Center. We particularly thank Mila Ilić and Marco Novais-Cruz for supporting characterization of cell lines and feedback on the manuscript. L.F. was supported by the European Union’s Horizon Europe research and innovation program under the Marie Skłodowska-Curie grant agreement No. 101152080. This work was further supported by Oncode Institute, which is partly supported by the Dutch Cancer Society (KWF).

---

**Figure S1.**
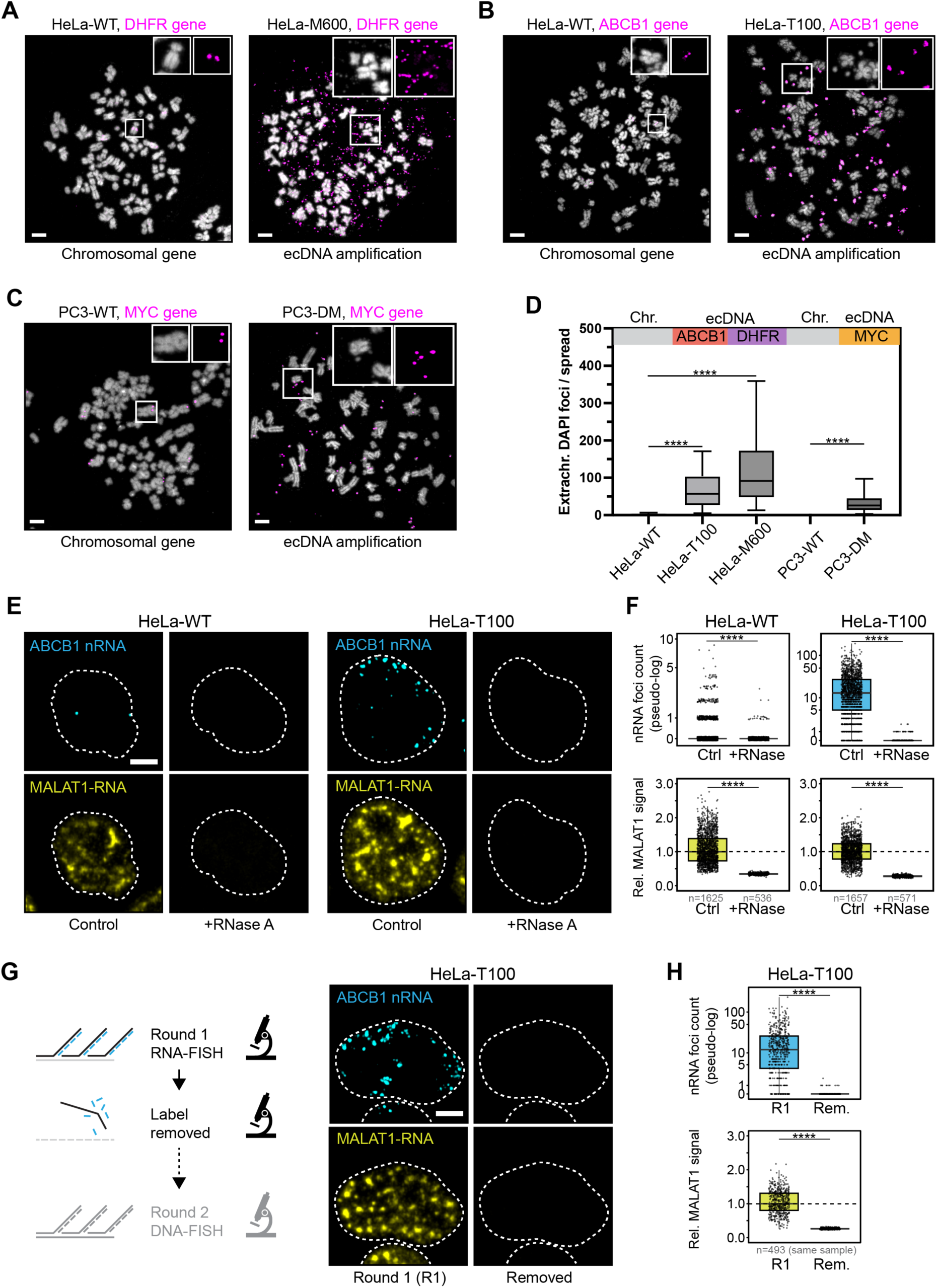
Characterization of ecDNA cell models and FLAGx assay specificity. (A-C) DNA-FISH on mitotic chromosome spreads confirming extrachromosomal amplification of *ABCB1*, *DHFR*, and *MYC* in ecDNA models compared with matched WT cell lines. (D) Quantification of extrachromosomal DNA foci on mitotic chromosome spreads (*n* = 60-119 quantified mitotic spreads per model across *N* = 3 independent experiments). (E, F) RNase A pretreatment efficiently depletes *ABCB1* nascent RNA (foci count per nucleus) and *MALAT1* RNA-FISH (total signal per nucleus) in both HeLa-WT and HeLa-T100 cells. (G-H) Validation of RNA-FISH label removal before the subsequent DNA-FISH round. Procedural details are outlined in the *Methods* section. Statistics: pairwise Wilcoxon rank-sum tests. All scale bars, 5 µm; *n*, number of cells.

**Figure S2.**
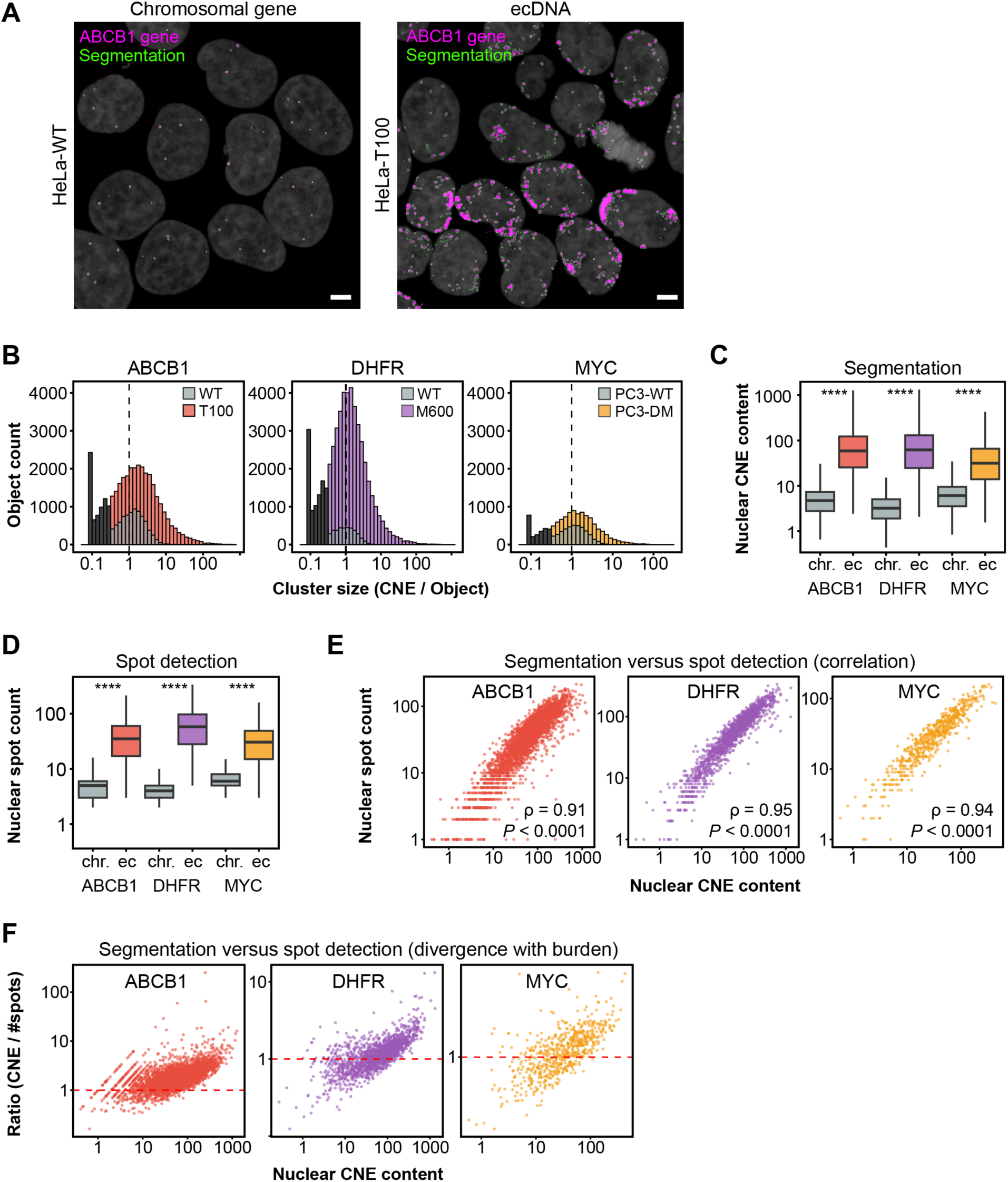
Validation of image-based ecDNA copy number quantification. (A) Representative DNA-FISH images showing gene object segmentation in HeLa-WT and HeLa-T100 (ec*ABCB1*) cells, used for CNE quantification. Scale bars, 5 µm. (B) Distributions of copy number equivalents (CNE) per gene object in matched WT (light gray) and ecDNA (colored) models. Dashed line marks CNE = 1; objects with CNE < 0.3 (dark gray) were excluded as background based on visual inspection. (C, D) Per-nucleus gene copy number estimated by segmentation-based CNE quantification (C) or segmentation-free *Spotiflow* spot detection (D) in matched chromosomal and ecDNA models. Asterisks indicate within-gene, pairwise Wilcoxon rank-sum tests comparing WT and ecDNA models (*P* < 0.0001). (E) Spearman correlation analysis for nuclear CNE content and *Spotiflow* spot counts, with correlation coefficients and *P* values indicated. (F) Ratio of nuclear CNE content to *Spotiflow* spot counts versus total nuclear CNE. Ratios >1 indicate higher copy number estimates by CNE quantification, particularly at high ecDNA burden per cell. Data across panels comprise all datasets from this study pooled (excluding drug treated and RNase-treated conditions): *N* = 2-7 independent experiments and *n* = 604-3178 cells per (gene x gene state) condition.

**Figure S3.**
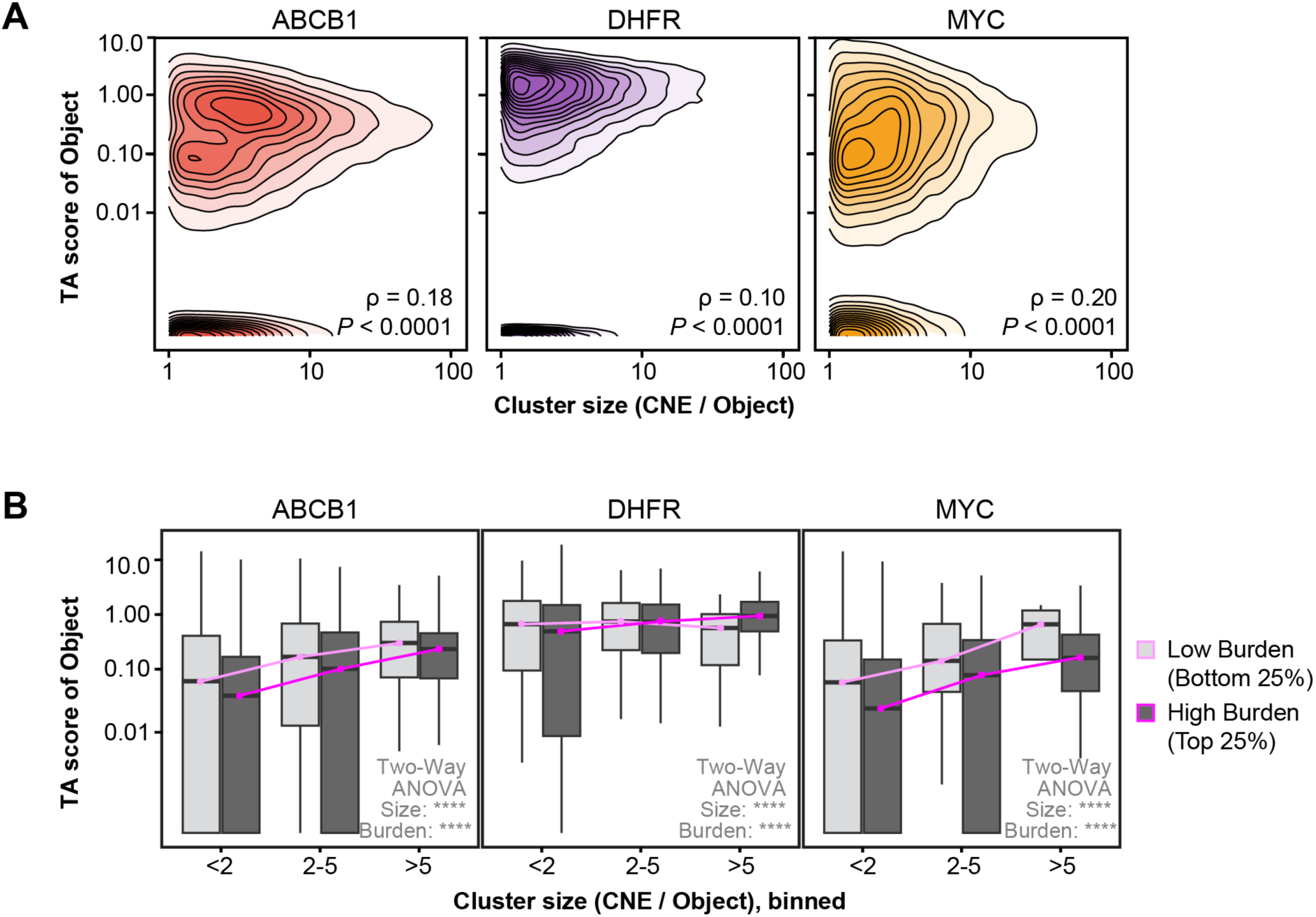
ecDNA clustering and copy number associations with transcriptional activity. (A) Correlation between ecDNA cluster size and per-copy transcriptional activity (TA), as in Figure 3E, excluding objects near the detection limit (CNE < 1). Spearman correlation coefficients (ρ) are indicated (*P* < 0.0001 for all genes). (B) Per-copy TA across three cluster size bins in cells with low (bottom 25%) or high (top 25%) ecDNA copy number burden. Lines connect group medians as guide to the eye. Two-way ANOVA (cluster size × nuclear CNE burden) for each gene: both main effects, *P* < 0.0001; interactions: ec*ABCB1*, *P* = 0.064; ec*DHFR*, *P* = 0.002; ec*MYC*, *P* = 0.714. Data across panels is identical to Figure 3.

**Figure S4.**
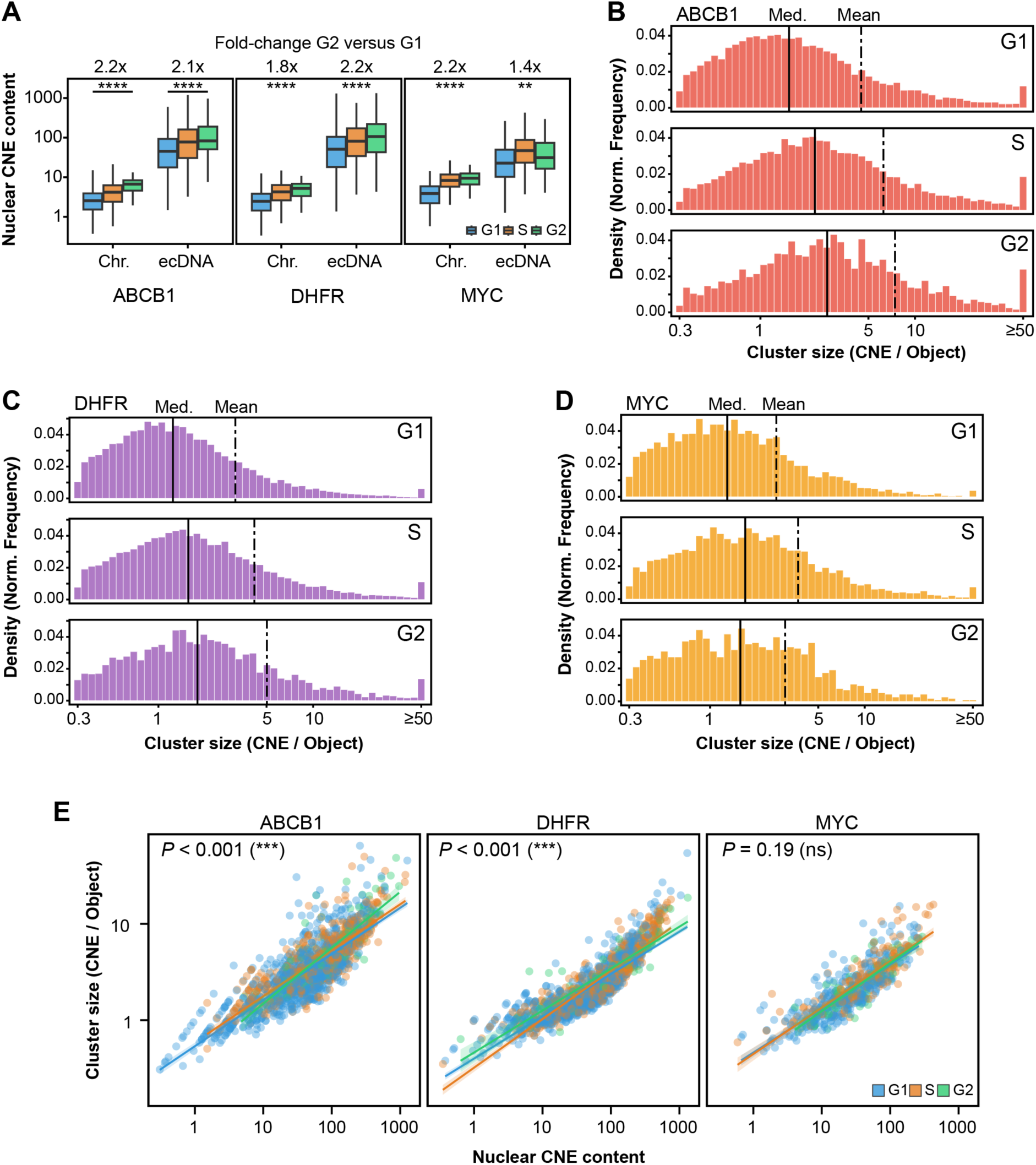
ecDNA copy number and clustering across the cell cycle. (A) Per-nucleus gene copy number (CNE) across G1, S, and G2 in matched chromosomal and ecDNA models of *ABCB1*, *DHFR*, and *MYC*. Copy number increased from G1 to G2 (Wilcoxon rank-sum tests, *P* < 0.01 for all genes). (B-D) ecDNA cluster size distributions across G1, S, and G2 for ec*ABCB1* (B), ec*DHFR* (C), and ec*MYC* (D) models. Cluster sizes increased from G1 to G2 (Wilcoxon rank-sum tests, *P* < 0.001 for all genes). (E) ANCOVA assessing the effects of nuclear CNE burden and cell cycle phase on ecDNA cluster size. Nuclear burden was significant for all genes (*P* < 0.0001); after adjusting for burden, cell cycle effects remained for ec*ABCB1* and ec*DHFR* (*P* < 0.001), but not ec*MYC* (*P* = 0.19). Data across panels is identical to in Figure 4, panels C-E.

**Figure S5.**
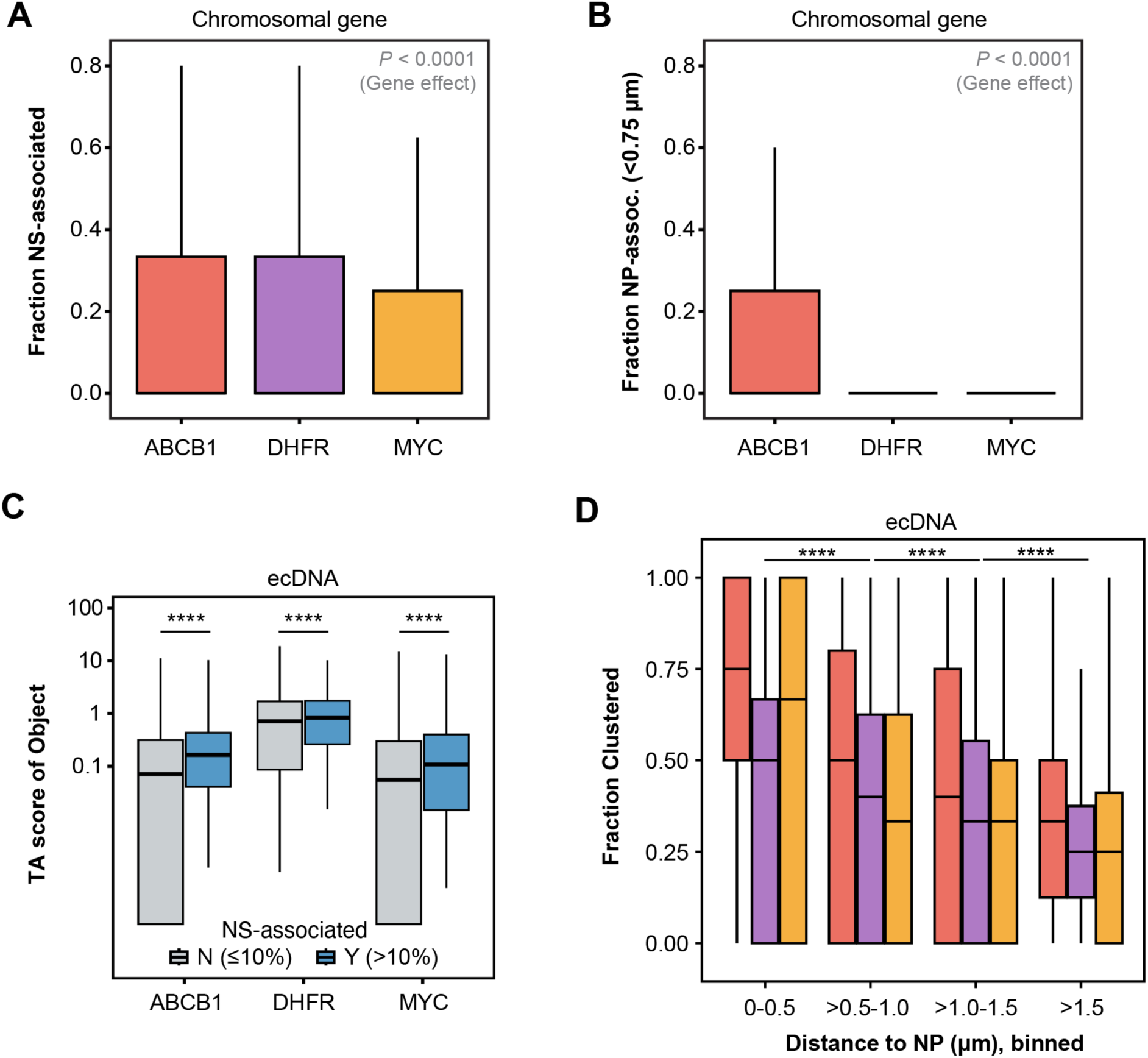
Chromosomal and ecDNA spatial dependencies. (A, B) Fraction of nuclear speckle (NS)-associated (A) and nuclear periphery (NP)-associated (B) chromosomal gene loci per nucleus, grouped by gene. One-way ANOVA: gene effects, *P* < 0.0001 for both (C) Per-copy transcriptional activity (TA) of ecDNA objects grouped by NS association. Tukey HSD, *P* < 0.0001 between associated and non-associated objects. (D) Fraction of clustered ecDNA per nucleus across NP distance bins, grouped by gene. Tukey HSD between distance bins, *P* < 0.0001. Data across panels is identical to Figure 5.

**Figure S6.**
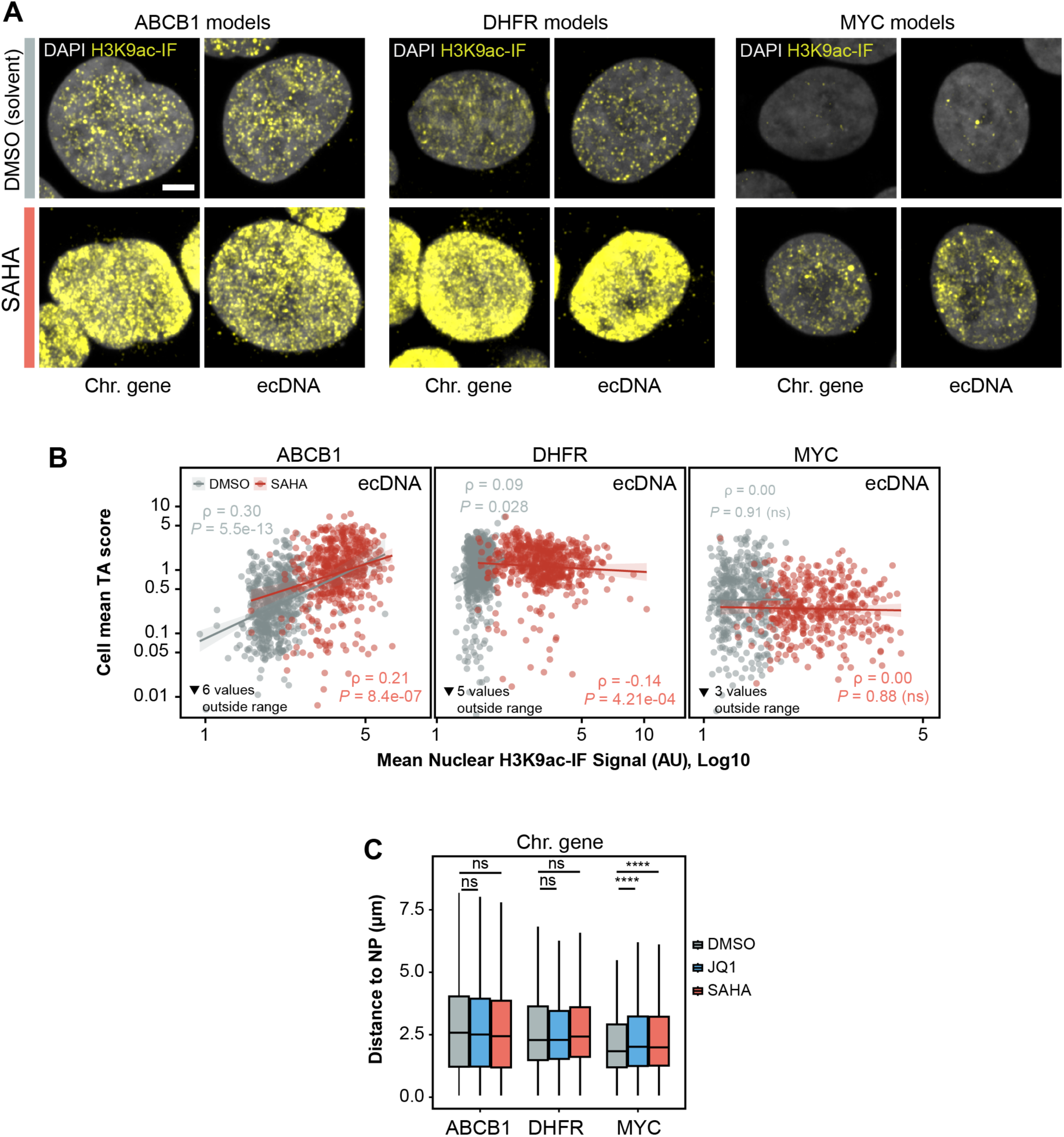
Histone acetylation and chromosomal gene positioning following SAHA treatment. (A) Representative FLAGx images of matched chromosomal and ecDNA models treated with SAHA (2 µM, 4 h) or DMSO, showing H3K9ac immunofluorescence (IF). Cells are identical to those shown in Figure 6A. Scale bars, 5 µm. (B) Correlation between mean nuclear H3K9ac IF intensity and mean per-copy transcriptional activity (TA) per nucleus for ec*ABCB1*, ec*DHFR*, and ec*MYC*. Each dot represents one cell. Spearman correlations were calculated separately for DMSO and SAHA treatments; coefficients (ρ) and *P* values are indicated. (C) Nuclear periphery (NP) distances of chromosomal *ABCB1*, *DHFR*, and *MYC* loci following JQ1 or SAHA treatment. Chromosomal *MYC* shifted modestly toward the nuclear interior after both treatments (Tukey HSD versus DMSO, *P* < 0.001). Data across panels is identical to Figure 6.

## METHODS

### Cell Culture

Cells were cultured at 37°C in 5% CO_2_ in Advanced DMEM/F12 medium (Gibco, 12634010) supplemented with 20 mM GlutaMAX (Gibco, 35050-038) and 50 U/ml Penicillin, 50 µg/ml Streptomycin (Merck, P4458) and 10% (v/v) Fetal Bovine Serum (Capricorn Scientific). For maintenance, cells were split every three to four days using Trypsin-EDTA/PBS solution (Gibco, 15400054) for detachment. Paclitaxel (Taxol), and Methotrexate (MTX) resistant HeLa cells with ec*ABCB1* (HeLa-T100) or ec*DHFR* amplification (HeLa-M600) were continuously grown in complete medium supplemented with 100 nM Taxol (Sigma-Aldrich, T1912) or 600 nM MTX (Sigma-Aldrich, A6770). HeLa-WT, PC3-WT and PC3-DM (ec*MYC* amplification) cells were cultured in complete medium without drugs. For assays, drug resistant cell lines were seeded in complete medium without drugs and cultured for at least 24 hours. All cell lines were regularly tested for the presence of mycoplasma using the MycoAlert mycoplasma detection kit (Lonza, LT07-318). Drug resistant ecDNA models were generate in-house.

### Cell seeding and treatment

For FLAGx assays, cells were seeded in either 8-well Nunc Lab-Tek II chambered cover glass vessels (Thermo Fisher, 155409PK) or in µ-Slide 8 Well high chambers (Ibidi, 80806) one to three days prior to staining at a density of 4-8×10^5^ cells per 500 µl per well, or 0.9-1.8×10^5^ cells per 500 µl per well, respectively. For EdU labeling, cells were treated with 10 µM EdU (Thermo Fisher, A10044) in preconditioned medium at 37°C in 5% CO_2_ for 15 minutes directly prior to fixation. For drug treatment experiments, cells were treated in preconditioned medium with either 0.2% (v/v) DMSO (Sigma-Aldrich, 472301) as solvent control, 2 µM SAHA (Abcam, ab144480), or 2 µM JQ1 (Abcam, ab146612) for 4 h at 37°C. Following treatment, cells were fixed and subjected to the FLAGx protocol described below.

### Oligonucleotides probes

FLAGx FISH probe pools tiling intronic and exonic regions of the target genes were designed using the OligoMinerApp^42^ with default parameters and a minimal binding temperature of 42°C. Probe targeting sequences were manually curated by BLAT search (UCSC genome browser) against human genome version GRCh38/hg38. Only probes with maximum one offtarget binding site were considered. Probe pools target the human genes *ABCB1* (133 intronic probes, NCBI gene ID 5243), *MYC* (50 intronic and 55 exonic probes, NCBI gene ID 4609), *DHFR* (112 intronic probes, NCBI gene ID 1719) and *MALAT1* (48 exonic probes, NCBI gene ID 378938). Telomere FISH probes were designed against canonical human TTAGGG_n_ repeats. Barcoded amplifier tails containing 20 nucleotide (nt) detection sequences (d1, d2, etc.) were subsequently added to the 3′ end of each targeting sequence. Targeting and detection barcode sequences were separated by a TT spacer according to the general probe structure: 5’-(28-41 nt unique targeting region)-TT-(20 nt barcode)4-3’. Secondary probes binding to the barcode sequences were conjugated to either 5’ATTO565 or 5’AlexaFluor647 fluorescent dyes. All probes are described in **Supplementary Table S1**. Oligonucleotides were manufactured by Integrated DNA Technologies (IDT Europe, Belgium).

### FLAGx staining

Incubations and washes were performed at room temperature (RT) using 250 µl of solution per well on adherent cells grown in cover glass bottom 8-well Nunc Lab-Tek II or µ-Slide 8 well high chambers, unless otherwise stated. All reagents used were RNA-grade and steps were carried out in an RNase-free environment.

For the RNA detection round (FLAGx round R1), the cells were first rinsed twice with 1×PBS (Thermo Fisher Scientific, AM9625). Cells were fixed for 15 minutes with 4%(w/v) paraformaldehyde (PFA) (EMS, 15735-10S) in PBS, followed by two PBS rinses. Cells were then permeabilized for 15 minutes with 100 µL of 0.5% (v/v) Triton X-100 (Sigma-Aldrich, T9284-500ML) in PBS, followed by two rinses in PBS. For RNase-treatment control experiments, permeabilization buffer was supplemented with 100 µg/ml RNase A (Thermo Fisher, EN0531). Following permeabilization, cells were equilibrated in 2×saline-sodium citrate (2×SSC) buffer (Thermo Fisher, AM9770) for 5 minutes. For primary probe hybridization, probes were pre-diluted in 80% (v/v) formamide (Merck, 344206) in nuclease-free water, supplemented with 1 µg/µl yeast tRNA (Invitrogen, AM7119). Probe pre-dilutions were then mixed with an equal volume of hybridization buffer, consisting of 30% (w/v) dextran sulfate (Merck, 265152-M), 2 mg/ml UltraPure BSA (Invitrogen, AM2616) in 4×SSC supplemented with 10 mM Vanadyl Ribonucleoside Complex/VRC (New England Biolabs, S1402S) RNase inhibitor. Probe concentrations used in the final hybridization mixture were 25 nM for *ABCB1* and *MALAT1*, 5 nM for *MYC*, and 10 nM for *DHFR* in a total hybridization volume of 200 µl per well. Next, sample chambers were sealed with Parafilm, placed in a humidified chamber, and incubated at 37°C for at least 16 hours or overnight. Following hybridization, samples were washed twice for 15 minutes each with 2×SSC/50% (v/v) formamide. Next, cells were rinsed once with 2×SSC and rinsed twice with PBS. For EdU-treated samples, incorporated EdU was fluorescently labeled with AlexaFluor 488 azide. In brief, the Click-iT EdU reaction was performed in 0.9 mM CuSO₄, 90 mM Tris-HCl (pH 8.5), 1.8% (w/v) ascorbic acid (Sigma-Aldrich, A92902) and 5 µM Alexa Fluor 488 azide (Thermo Fisher, A10266) for 15 minutes in the dark. Stained samples were then rinsed twice with 500 µl of 2×SSC. For samples not treated with EdU, cells were equilibrated directly in 2×SSC for 5 minutes following the post-hybridization washes. For both EdU-treated and untreated samples, secondary hybridization was performed using 10 nM fluorescently-conjugated detection probe in 10% (v/v) formamide/2×SSC for at least one hour at 37°C in a humidified chamber. Cells were subsequently incubated with 5 µM DAPI (Sigma-Aldrich, D9542) in 2×SSC for 5 minutes, rinsed three times in 2×SSC, and then stored in 4×SSC buffer supplemented with 5 mM VRC RNase inhibitor at 4°C in the dark until imaging.

For the DNA detection round (FLAGx round R2), storage buffer was removed and the cells were rinsed twice with PBS and permeabilized for 15 minutes in 0.5% (v/v) Triton X-100 in PBS supplemented with 100 µg/ml RNase A (Thermo Fisher, EN0531), followed by two rinses with PBS. Samples were treated with 0.1 M hydrochloric acid for 15 minutes, neutralized in 2×SSC buffer for 5 minutes, and equilibrated in 2×SSC/50% (v/v) formamide for 30 minutes. For primary probe hybridization, probes were pre-diluted in 85% (v/v) formamide (Merck, 344206) in nuclease-free water, supplemented with 0.2 µg/µl yeast tRNA (Invitrogen, AM7119) and 0.05 µg/µl human Cot-1 DNA (Invitrogen, 15279011). Probe pre-dilutions were then mixed with an equal volume of hybridization buffer as described for the RNA detection round. Probe concentrations used in the final hybridization mixture were 10 nM for *ABCB1*, *MALAT1* and telomeres, 5 nM for *MYC*, and 3.3 nM for *DHFR* in a total hybridization volume of 200 µl per well. After addition of the hybridization solution, samples were sealed, heat-denatured at 80°C for 7 minutes on a heat block surface and subsequently incubated at 37°C for at least 16 hours or overnight in a humidified chamber. Following primary probe hybridization, samples were washed twice with 2×SSC for 5 minutes each to remove residual hybridization solution. Afterwards, samples were washed three times in 1×SSC buffer and three times in 0.1×SSC buffer for 5 minutes each at 40-42°C on a heat block surface. Samples were then equilibrated for 5 minutes in 2×SSC buffer at RT. Secondary probe hybridization, subsequent washes and storage without RNAse inhibitor were performed as described for round R1. For FLAGx destaining control experiments, the full RNA detection round was performed and imaged. However, to detect residual RNA signal carryover, samples were subjected to an additional round of imaging after primary DNA probe hybridization and the 40-42°C washes, but prior to secondary DNA probe hybridization.

### FLAGx with immuno-staining

For samples subjected to combined FLAGx and immunofluorescence staining (IF), the RNA detection protocol was followed as described above up to and including the post-hybridization washes. Next, cells were blocked for at least 30 minutes at RT in 100 µl of 1 mg/ml UltraPure BSA in PBS-0.1%(v/v) Tween-20 (PBST) supplemented with 5 mM VRC (IF blocking buffer).

Next, the blocking buffer was removed and samples were incubated in 100 µl per well of primary antibody solution containing rat anti-H3K9ac antibody (Active Motif, 61664) diluted 1:800 in IF blocking buffer overnight at 4°C in a humidified chamber. The following day, cells were washed three times for 5 minutes each in PBST at RT. Next, the secondary probe hybridization was carried out as described above, but with the addition of goat-anti-rat AlexaFluor488 secondary antibody (Thermo Fisher, A-11006) diluted 1:500 in the secondary hybridization mix supplemented with 0.4 mg/ml UltraPure BSA. Subsequent washes, DAPI staining and sample storage were carried out as described above.

### Mitotic chromosome spreads and DNA-FISH validation

Cells grown to approximately 70% confluency in a T75 culture flask, were treated with 0.125 µg/ml Nocodazole (Sigma-Aldrich, M1404) in fresh medium for 6 to 16 hours to induce mitotic arrest. Mitotic cells were collected by mechanical shake-off and subsequent centrifugation for 5 minutes at 1500 rpm at RT. Cell pellets were resuspended in 1 ml of pre-warmed (37°C) 75 mM potassium chloride solution dissolved in water and incubated for 10 minutes at 37°C. Five medium-sized drops of fixative (3 methanol : 1 acetic acid (v/v)) were added to the cell suspension, followed by centrifugation for 5 minutes at 1500 rpm. The cell pellet was resuspended in 1 ml of fixative and incubated for 15 minutes at RT. The fixative was subsequently replaced once by centrifugation, and the cells were incubated in fresh fixative for an additional 15 minutes. Fixed cells were resuspended in fixative containing 2.8 µM DAPI and stored at -20°C until use. For preparation of mitotic chromosome spreads, fixed and DAPI-stained cell suspensions were dropped on standard microscopy glass slide or into 8-well cover glass bottom chambers from a height of 10-15 centimeters. The samples were allowed to airdry until the fixative had evaporated. Slides were mounted with Prolong Gold (Invitrogen, P36934) using 25x50 mm cover glasses. For DNA-FISH staining of target genes on mitotic chromosome spreads, samples dispensed in 8-well cover glass bottom chambers were dried for at least 30 minutes at RT, then again fixed in 4% (w/v) PFA/PBS for 5 minutes at RT. Following fixation, mitotic spreads were processed using the DNA-FISH FLAGx R2 staining protocol described above.

### Image acquisition

Samples were imaged on a Leica Stellaris 8 Falcon confocal system (Leica Microsystems) with 8 kHz resonant scanner, equipped with a 63x/1.40 oil immersion objective and HyD S/X/R detectors. A tunable range 440-790 nm White Light Laser (WLL) and 405 nm diode laser were used for excitation. DAPI, AlexaFluor488, ATTO565, and AlexaFluor647 fluorescence signals were detected as follows in two sequential scans (DAPI/ATTO656; AF488/AF647): DAPI (ex: 405 nm, em: 430-494 nm), AlexaFluor488 (ex: 499 nm, em: 510-570 nm), ATTO565 (ex: 575 nm, em: 600-650 nm), AlexaFluor647 (ex: 652 nm, em: 663-750 nm). Nascent RNA and DNA signals of the target genes were detected in the ATTO565 channel, IF and EdU in AF488 and MALAT1, telomeres in AF647. Images were acquired in 16-bit format with field of view (FOV) dimensions of 2048 × 2048 pixels (pixel size: 70 nm), using a scanning speed of 700 Hz, line averaging of 2 and unidirectional scanning. For FLAGx samples, selected regions of ∼400x400 µm in xy and 12 µm in z (step size 0.4 µm) were recorded as 3x3 tile scans (10% tile overlap). One to two regions were acquired per condition and independent FLAGx experiment. After the RNA detection round, positional information of the regions was documented and the same regions were scanned with identical acquisition settings in the subsequent DNA detection round. Region tile scans were stitched into coherent image regions and then maximum-intensity projected using the Leica LASX software. Images of mitotic spreads stained by DAPI or mitotic spreads co-stained by DNA-FISH were acquired with the same instrument setup.

### Image preprocessing

Stitched and projected image regions of the matched RNA and DNA FLAGx rounds were registered using the Register Virtual Stack Slices FIJI^43^ plugin with an elastic registration algorithm (bUnwarpJ^44^). DAPI channels of both rounds were used for alignment using the RNA round as reference. After registration, all other channels were xy-transformed accordingly using the Transform Virtual Stack Slices FIJI plugin. The resulting image regions were cropped to remove alignment artifacts at the image region borders.

### Image analysis

Cell nuclei were segmented on DAPI images of both rounds using Cellpose^45^ v3.0.12 with the pre-trained model ‘nuclei’ and diameter 200. For segmentation-free and threshold-agnostic FISH spot detection, Spotiflow^33^ v0.5.0 was used with the pre-trained model ‘general’. Segmentation of gene objects (chromosomal loci and ecDNA) in the DNA-FISH channel and nuclear speckles in the MALAT1 channel followed a two-step procedure. First, images were subjected to a segmentation parameter sweep entailing different strengths of Gaussian blurring followed by per nucleus median intensity-based pixel thresholding at different stringencies (factor above median). Second, the optimal parameter combination for each target channel was identified by visual inspection. Identical segmentation parameters were then applied for images of the same gene target. Object segmentation and all downstream image processing including intensity feature extraction, assignment to nuclei and QC and was implemented using custom workflows in R (https://www.R-project.org/) and functions based on the EBImage R package^46^, mostly as described previously^47,48^. The main custom image processing and feature extraction workflow used in this study are documented in the CellAnalyzeR R package (doi.org/10.5281/zenodo.23659910) and associated image processing script collection (doi.org/10.5281/zenodo.23660971).

Segmented nuclei and objects were imported as semantic pixel masks and intensity features in these image regions were measured across all relevant channels. Subnuclear objects were mapped to nuclei based on spatial overlap analysis. Overlap of objects with other object types, such as nuclear speckles, were scored accordingly. For spots detected using Spotiflow^33^, the native spot coordinate output was converted into single pixel semantic masks representing spot centers and processed as the object masks above for assignment to individual nuclei. To determine the distance of subnuclear objects to the nuclear periphery, shape and position features were computed for each object. Then, the shortest distance to the nuclear mask edge was calculated using a Euclidean distance transform map (*distmap* function) derived from the parent nucleus mask. For binary spatial classification, objects were defined as nuclear periphery (NP)-associated if their center of mass was ≤0.75 µm from the nuclear edge, and nuclear speckle (NS)-associated if ≥10% of their pixel area overlapped with a segmented MALAT1 mask.

To quantify transcription, the integrated pixel intensity (sum) was determined in both the DNA-FISH and RNA-FISH channel for each gene object mask alongside with the background signal of the parent nucleus (mean intensity outside objects masks). After channel specific background subtraction, the DNA to RNA signal ratio was determined as raw transcriptional activity (TA) value. Objects with a negative raw TA score after background subtraction procedures were floored to a pseudo value of 0.0001. Per object TA values then were transformed into per-copy TA scores using the conversion factor of integrated DNA-FISH pixel intensity to copy number equivalent (CNE), which was determined for each gene target individually. Copy number equivalents were defined in two steps. First, background-corrected sum DNA-FISH signal intensities were determined for each segmented object across all WT datasets from one gene condition (HeLa-WT for *ABCB1* and *DHFR*; PC3 for *MYC*). Per gene target, CNE of a single gene copy (singlet) was anchored to 1 and to the median of DNA-FISH sum intensity values of objects in unamplified WT cells that ranged between of 30-60 pixels in area. This threshold was established by visual inspection. Subsequently, all object DNA intensity sum values in WT and ecDNA models were normalized to this gene-specific anchor value to obtain a CNE value for each object. Subnuclear gene objects were further classified as singlets (CNE < 2.0) or clusters (CNE ≥ 2.0). To determine the total ecDNA copy number threshold (T_Clust_) required to drive 25% of ecDNA into clusters, a logistic regression model was fit to the single-nucleus clustering fractions as a function of total nuclear CNE content. Total CNE content was defined as the sum of CNE values of all valid objects within that nucleus.

For all analyses presented, gene objects with a CNE value ≤ 0.3 were excluded as segmentation noise. This threshold was established based on visual validation. The fraction of transcriptionally active objects was calculated independently of CNE values and the TA scores using a fixed measure of RNA signal contrast in the pixels corresponing segmented objects regions. Objects were defined as transcriptionally active if their mean nascent RNA signal intensity exceeded the mean local nuclear background by a factor of 1.2-fold or more. To assess intra-nuclear transcriptional variability, the coefficient of variation (CV) of the per-copy TA scores was calculated for all valid gene objects within a single nucleus. Nuclei with a single segmented object were excluded from CV analysis, since the CV could not be determined for these cases.

### Analysis of cell cycle and mitotic cells

For cell cycle stratification of FLAGx datasets, cells were gated based on the integrated DAPI signal intensity (sum) per nucleus used as measure DNA content and the nuclear mean EdU signal as indicator of S-phase. Per-image region adaptive thresholding was used to account for technical variance. S-phase cells were identified by an EdU mean intensity exceeding a dynamic threshold, defined as the sample median EdU intensity plus 30% of the difference between the 95th percentile and the median. To establish the G1/G2 DNA content boundary, the primary G1 peak was identified via density estimation of the DAPI sum signal, and the G2 threshold was set at 1.5-fold of this G1 peak. Cells exceeding this threshold and negative for EdU were classified as G2, while the remaining double-negative cells were classified as G1.

FLAGx-stained mitotic cells were identified by their nuclear morphology in the DAPI channel by scoring on fully preprocessed and aligned FLAGx datasets. Image regions containing individual mitotic cells were isolated by manual cropping. The resulting crops were then processed using the analysis workflows described for regular FLAGx regions above. However, each mitotic cell was manually curated by visual inspection post-processing to flag cells with unsucessful segmentation and exclude cells with less than two discernable gene objects. The number of ecDNA foci on images of DAPI-stained mitotic chromosome spreads used for cell model validation was quantified by manual counting in FIJI.

### Image representation

Unless stated otherwise, all microscopy images shown in figures are cropped maximum-intensity z-projections of the raw image stacks. To enhance visibility, Gaussian smoothing (sigma 1.5) was applied to all images presented, alongside linear intensity range adjustments to enhance contrast for optimal visibility. Per channel and target, identical contrast adjustments were applied within multi-image panels and across figures, where possible.

### Quantification and statistical analysis

Quantification and statistical analyses were performed in R v4.4.0. In figure legends, *N* denotes the number of independent biological experiments, and *n* denotes the number of analyzed single cells, nuclei, or gene objects per condition. Two-group comparisons were evaluated using pairwise Wilcoxon rank-sum tests. Multi-group and multi-factorial continuous data (e.g., spatial bins, cell cycle phases, and gene states) were analyzed using One-Way or Two-Way analysis of variance (ANOVA), followed by Tukey’s Honest Significant Difference (HSD) post-hoc tests to control the family-wise error rate during multiple comparisons. Analysis of Covariance (ANCOVA) was used to evaluate group effects while controlling for continuous covariates, such as total ecDNA copy number. Significance is indicated as * *P* < 0.05, *P* < 0.01, *** *P* < 0.001, and **** *P* < 0.0001; ns, not significant. Boxplots depict the median and interquartile range (IQR), with whiskers extending to the most extreme data points within 1.5 × IQR of the first and third quartiles.

### Declaration of generative AI and AI-assisted technologies in the manuscript preparation process

During the preparation of this work, the authors used ChatGPT to check grammar and refine selected text passages. They further used ChatGPT and Google Gemini to assist development of R code for image analysis, graph plotting and associated statistical testing (within R). After using these tools, the authors reviewed and edited the content and/or code as needed and take full responsibility for the content of the publication.

